# Condensate-based shells and scaffolds via interfacial liquid-to-solid transition of disordered peptides

**DOI:** 10.1101/2025.09.10.674898

**Authors:** Chang Chen, Qimeng Wang, Nele Jarnot, Ingrid Dijkgraaf, Siddharth Deshpande

## Abstract

Living organisms use biomolecular condensates to respond to dynamic environments and create functional materials with complex architectures. Exploring such phase-separated systems beyond the naturally occurring scenarios may offer valuable insights for emergent synthetic biosystems. Here, we report the self-assembly behavior of a short, disordered peptide sequence (termed CT45) derived from a protein present in the bioadhesive system of tick ectoparasites. We show that CT45 spontaneously accumulates at polar-nonpolar interfaces, and further undergoes liquid-liquid and liquid-to-solid phase transitions to create mechanically stable structures. When encapsulated within vesicles and presented with a stable oil-water interface, CT45 rapidly forms solid shells, which can be reinforced by up-concentrating the material through osmotic imbalance. Unexpectedly, when presented with a transient acetone-water interface, CT45 condenses at the evaporating interface and forms interconnected, porous mesoscopic scaffolds. The underlying mechanism is found to be the amphiphilic nature of CT45 leading to interfacial accumulation, enhancing intermolecular *π*-based interactions to trigger phase transitions. The micron-sized shells exhibit appreciable mechanical strength and the porous scaffolds present a highly stable platform capable of retaining molecules. In conclusion, the presented condensate-based microscopic and mesoscopic scaffolds hold significance in customizable condensate architectures, with potential applications in biomedical engineering and synthetic biology.

## Introduction

Over the course of evolution, organisms have developed sophisticated strategies to adapt to dynamic environmental changes by utilizing basic molecular building blocks.^1^ Biomolecular condensates serve as a crucial mechanism for reorganizing molecular components and endowing them with functional properties in response to environmental stimuli.^2–4^ In recent years, increasing attention has been directed toward understanding the behavior of condensates within cells, with the aim of elucidating their role in cellular functions and replicating these life-like behaviors in artificial systems.^5–10^ However, beyond the confined environments of cellular compartments, the condensation and solidification behaviors at extracellular level has also received significant attention. ^11–13^ Such liquid-liquid phase separation (LLPS) behaviors may represent either adaptive responses to environmental fluctuations or active, purpose-driven strategies employed by living systems.

The attachment of a tick’s mouthparts to the host skin during blood feeding showcases a recent example of a secretory process where LLPS and biocondensation might be involved. ^11^ Ticks harbor glycine-rich proteins (GRPs) in their saliva, which undergoes liquid-to-solid transition to form a cement cone around the incision site. ^14^ A high ratio of glycine residues makes GRPs conformationlly flexible, thereby promoting cation−*π* or *π* − *π* interactions between arginine and aromatic amino acid residues.^11^ Thus, GRPs hold significant potential as a basic unit for condensate architectures.

Environmental gradients are known to be an important means of controlling protein enrichment and condensation. For example, salt gradient induced by environmental salinity change induces internal secondary phase separation within preformed condensate, thus switching the glycinin condensates to a hollow structure.^15^ Solvent gradient can also be applied to assemble condensate capsules using peptides derived from the cuticle of *Ostrinia furnacalis*, owing to their intrinsic affinity for specific solvent concentrations. ^16^ However, in most cases, the fundamental basis for molecular affinity toward particular solvents or concentration regimes remains poorly understood. Although the mechanisms behind their assembly are not yet fully understood, condensate-based capsules have demonstrated superior performance compared to coacervate droplets, particularly in terms of kinetic stability, cargo loading and protection, cellular uptake efficiency, and enhanced therapeutic efficacy.^16,17^ Moreover, their mechanical properties offer exciting prospects for expanded practical applications.^17^ As independent units, condensate vesicles can also be regarded as the basic unit of artificial tissue-like constructs. ^18^ Accordingly, establishing high-throughput and stable fabrication protocal for condensate vesicles will be pivotal for advancing their practical applications, particularly in drug delivery or synthetic biological systems.

Here we describe an innovative strategy for designing mechanically robust peptide-based coacervate shells via polarity gradient-mediated self-assembly. Using an LLPS-prone peptide derived from a tick salivary protein as the self-assembling unit, we systematically analyzed its affinity behavior on polar-nonpolar interfaces and elucidated their interaction mechanism. Employing a microfluidic system, we generated double emulsion vesicles with an oil–water interface, which facilitated peptide accumulation at the interface, subsequent coacervation, and liquid-to-solid phase transition, forming micron-sized condensate shells. Osmolarity-driven interfacial crowding significantly enhanced peptide interactions and mechanical strength of the formed vesicles. Finally, supported by the transient polarity gradients in acetone-water system, we constructed peptide-based, porous mesoscopic scaffolds, spanning hundreds of micrometers. Such condensate shells and subsequent hierarchical assemblies offers a potential avenue in engineering biocondensate-based architectures, with broad applications in biomedical engineering and synthetic biology.

## Results

### Tick salivary peptide as the basis for building synthetic scaffolds

Based on our recent findings on the LLPS and liquid-to-solid transition of a tick salivary GRP,^11^ we decided to further test its potential in forming peptide-based shells and higher assemblies. GRP (UniProt Q4PME3) consists of a 19-amino acid-long signal peptide at the N-terminus (sequence, ^1^MNRMFVLAATLALVGMVFA^19^) and a 77-amino acid-long disordered domain. Particularly, we focused on the 45-amino acid-long C-terminus fragment (CT45; Figure 1a, structure predicted by AlphaFold^19^) as it was identified as the main promoter for LLPS. Being rich in arginine (R) and aromatic (F, Y) amino acid residues (in total 20 %, see Figure 1b), CT45 can form extensive cation−*π* and *π* − *π* interactions for rapid and robust condensation. ^11^ Among these residues essential for LLPS, 56% are polar (Y, R) and 44% are nonpolar (F). Meanwhile, LLPS-forming potential of CT45 is also reflected in the prediction by the AIUPred algorithm,^20^ with a score above 0.5 for the entire sequence (Figure 1c). All things considered, we synthesized CT45 using solid-phase peptide synthesis (see Methods for details) and assessed its potential in forming micro-scaffolds (Figure 1d).

**Figure 1:**
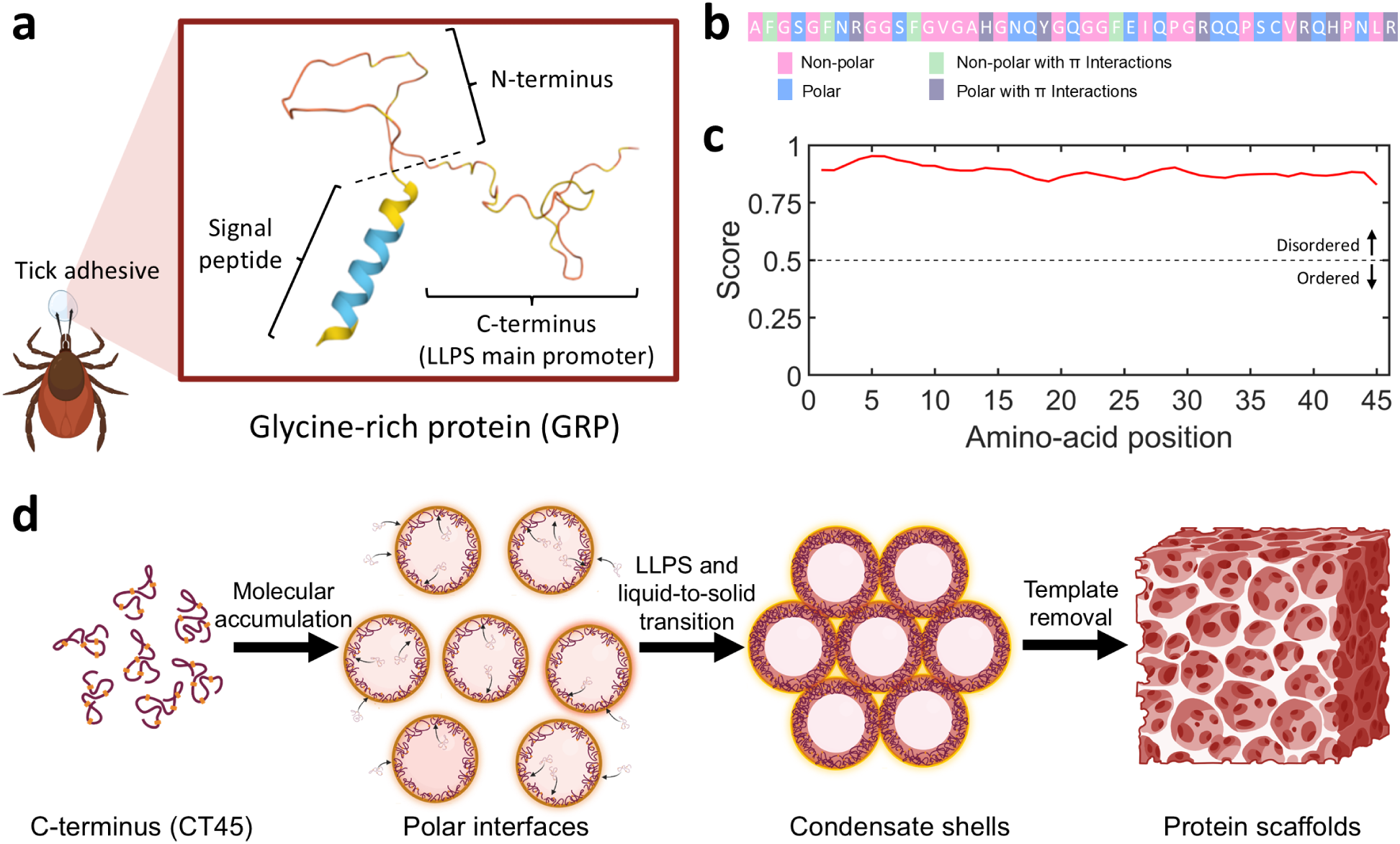
Design principle behind the self-assembly of the C-terminus peptide using emulsion templates to form solid shells and mesoscopic scaffolds utilizing solvent polarity gradients. **(a)** The C-terminus peptide (CT45) is derived from a tick salivary GRP, with a putative role in the tick cement cone formation. AlphaFold model correctly predicts an *α*-helical N-terminal signal peptide and a disordered structure for the rest of the sequence, with CT45 being the main inducer of LLPS. **(b)** Amino-acid composition of CT45 shows 53% of nonpolar amino acids and 24% residues capable of forming cation*−π* or *π − π* interactions. **(c)** The AIUPred scores of the entire CT45 sequence are above 0.5, indicating CT45 to be a highly disordered protein. **(d)** Conceptual sketch illustrating the formation process of emulsion-templated CT45-based scaffolds.

### Spontaneous CT45 accumulation at the oil-water interface

Given its strong inclination to undergo LLPS, we investigated whether CT45 could behave similarly when adsorbed on specific surfaces. Utilizing the Kyte-Doolittle hydropathy scale,^21^ CT45 can be seen as moderately hydrophilic as the average hydropathy score is ≈ –0.85 (Figure 2a, pink bar). Meanwhile, significant presence of arginine residues (9 %) imparts a net positive charge to CT45 at pH 7.4 (Figure 2a, blue bar). Therefore, we began with studying the interaction of CT45 to a hydrophilic, negatively charged surface. Using a lab-on-a-chip device, we flowed 125 *µ*M CT45 (CT45:carboxytetramethylrhodamine (TAMRA)-CT45 = 10:1, molar ratio; the latter was added for fluorescent visualization) in PBS at pH 7.4 into microwells coated with a lipid bilayer (Figure 2b, see Supplementary Figure 1a and Methods for details). The composition of the lipid bilayer was 89.9 % 18:1 (Δ9-Cis) PC (DOPC) incorporation of 10 % 18:1 PS (DOPS), imparting negative charge on the bilayer, and 0.1% 18:1 Cyanine 5 PE (cy5-DOPE) for fluorescent visualization. Figure 2c shows that the lipid bilayer were evenly formed as visually judged by a smooth, homogeneous surface, but more importantly, validated through a fluorescence recovery after photobleaching (FRAP) assay (Supplementary Figure 1b-c). However, even after 1 h of incubation, CT45 could be easily washed away when purged by a buffer solution. As indicated by Figure 2d, normalized TAMRA-CT45 intensity decreased from 0.92 to 0.08 immediately after flowing a CT45-free buffer. The final fluorescence intensity of near-zero (right inset in Figure 2d), clearly suggested that CT45 did not adsorb to the negatively charged lipid bilayer.

**Figure 2:**
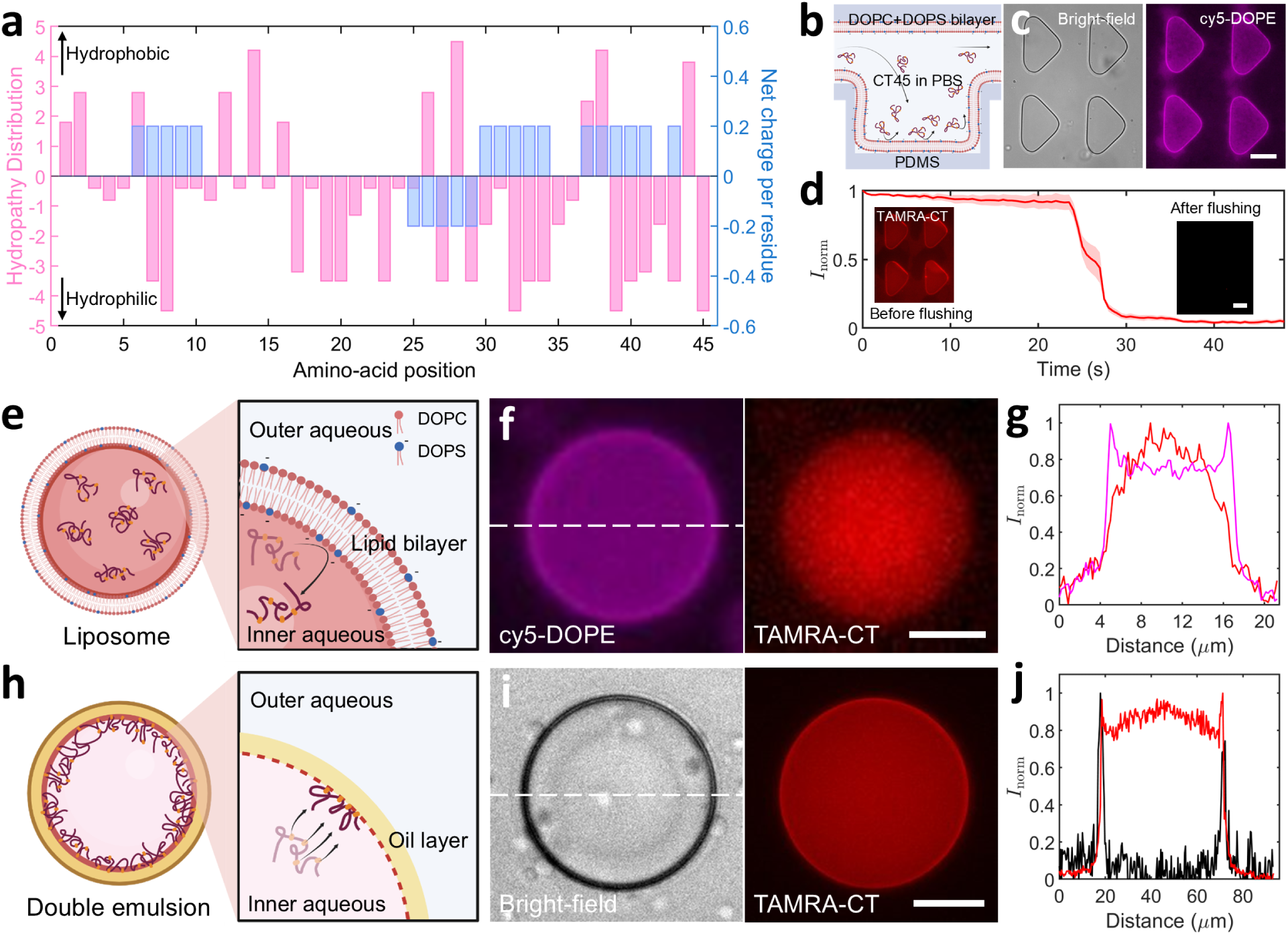
CT45 spontaneously accumulates at the oil-water interface. **(a)** Hydropathy distribution based on Kyte-Doolible hydropathy model (pink) and net charge per residue obtained using CIDER42 (blue), both plotted as a function of the CT45 sequence, showing that it is an amphipathic and moderately positively charged peptide. The net charge for each amino acid quantifies the average electrostatic charge of a 5-residue window centered on that amino acid. **(b)** Schematic diagram showing the side-view of a bilayer-coated microwell array. **(c)** Bright-field and fluorescence images show the top views of the microwell array covered with a phospholipid bilayer. Scale bar, 20 *µ*m.**(d)** A plot showing rapid reduction in the CT45 fluorescence within the microwells upon flushing the device with PBS; the insets illustrate fluorescence images before and after washing. The lipid bilayer composition was 89.9 % DOPC + 10 % DOPS + 0.1% cy5-DOPE (molar ratio), and 125 *µ*M CT45 (CT45:TAMRA-CT45 = 10:1, molar ratio) in PBS was incubated. Data points represent mean values while the shaded areas indicate standard deviations (n = 5 microwells within one microfluidic device). Scale bar, 20 *µ*m. **(e)** Conceptual sketch illustrating that CT45 molecules do not interact with the negatively charged membrane of the liposome. **(f)** Fluorescence images showing the equatorial plane of a liposome (left) and the encapsulated CT45 distribution (right). Scale bar, 5 *µ*m. **(g)** Normalized fluorescence intensity (magenta: lipids; red: CT45) corresponding to the dotted line in panel (f) showing a homogeneous CT45 distribution within the lumen. **(h)** Conceptual sketch illustrating that CT45 molecules accumulate at the oil-water interface offered by the double emulsion confinement. **(i)** Bright-field (left) and fluorescence (right) images showing the equatorial plane of a DE droplet and CT45 accumulation on its inner surface. Scale bar, 20 *µ*m. **(j)** Normalized gray value (black) and fluorescence (red) intensity corresponding to the dotted lines in panel (i) show accumulation of CT45 on the inner surface of the DE confinement. In e-j, the encapsulated mixture consisted of 55 *µ*M CT45 (CT45:TAMRA-CT45 = 10:1, molar ratio) and 300 mM Na_2_HPO_4_. In e-g, the lipid bilayer composition was 89.9 % DOPC + 10 % DOPS + 0.1% cy5-DOPE (molar ratio). In h-j, the oil phase was fluorinated oil (HFE 7500) containing 2% FluoSurf-C surfactant.

Given the indication that electrostatic interactions did not promote CT45 absorbance or were not strong enough to localize CT45 to the interface, we proceeded with enclosing CT45 within a fully confined hydrophilic interface and incubating it for an extended period to see if it changes the situation. Utilizing a well-established technology, octanol-assisted liposome assembly (OLA),^22–24^ we produced liposomes with the same lipid composition as the previous experiment in the microwells and encapsulated 55 *µ*M CT45 (CT45:TAMRA-CT45 = 10:1, molar ratio) and 300 mM Na_2_HPO_4_ in their lumen (Figure 2e). The idea behind encapsulating Na_2_HPO_4_ was to aid eventual CT45 phase separation,^11^ and this recipe was used in subsequent experiments. Even after 24 h of incubation, the encapsulated CT45 (Figure 2f, right) showed no detectable affinity for the hydrophilic surface (Figure 2f, left). As shown in Figure 2g, the TAMRA-CT45 intensity was predominantly distributed inside the lumen, rather than at the lipid bilayer.

Since the overall hydrophilicity (hydropathy *<* 1) and charge (net charge *>* 1) of CT45 does not lead to its adsorption at the interface, we turned our attention to its nonpolar parts, given the presence of 53% hydrophobic residues in the CT45 sequence (Figure 1b). To do so, we used another established biocompatible on-chip platform^25^ to encapsulate same CT solution as liposome experiment in water-in-oil-in-water double emulsion droplets (DEs). Figure 2h illustrates that the thin oil shell, functioning as the middle phase, establishes a hydrophobic surface in contact with the inner aqueous phase. In contrast to the lipid bilayer and liposoem experiments, we observed that the encapsulated CT45 molecules spontaneously adsorbed at the oil-water interface when encapsulated in DEs. As shown in Figure 2i, this recruitment of CT45 resulted in the formation of a fluorescent shell along the inner surface of the DE. The overlap between the TAMRA-CT45 fluorescence intensity peak and the bright-field grayscale profile (Figure 2j) further confirms this specific adsorption. Thus, we demonstrated the ability of CT45 molecules to adsorb at the hydrophilic-hydrophobic interface, likely due to the presence of nonpolar residues.

### CT45 undergoes liquid-to-solid transition at stable interfaces

The surface adsorption of CT45 molecules provided us with the possibility to investigate emulsion-templated CT45 condensation and possible subsequent structural transformations, as this interfacial adsorption may promote solidification of CT45 molecules. Figure 3a shows that nonpolar residues of CT45 preferentially coming into contact with the hydrophobic oil phase, and are designated as ‘coaters’. Due to this coating, the CT45 molecules are passively crowded and get concentrated at the 2D interface. CT45 is also rich in glycine (24%), which enhances the flexibility of the chain (thus being seen as a ‘spacer’). The presence of these spacer residues promotes intra-and intermolecular interactions of the CT45 molecules, during their accumulation at the surface at high concentrations. ^26^ This likely leads to the sticker residues forming cation−*π* and *π*−*π* interactions, culminating in condensation. Since GRP condensates undergo aging and form a solid-like state,^11^ we expected CT45 condensates to also eventually solidify after the crowding-induced condensation at the interface.

**Figure 3:**
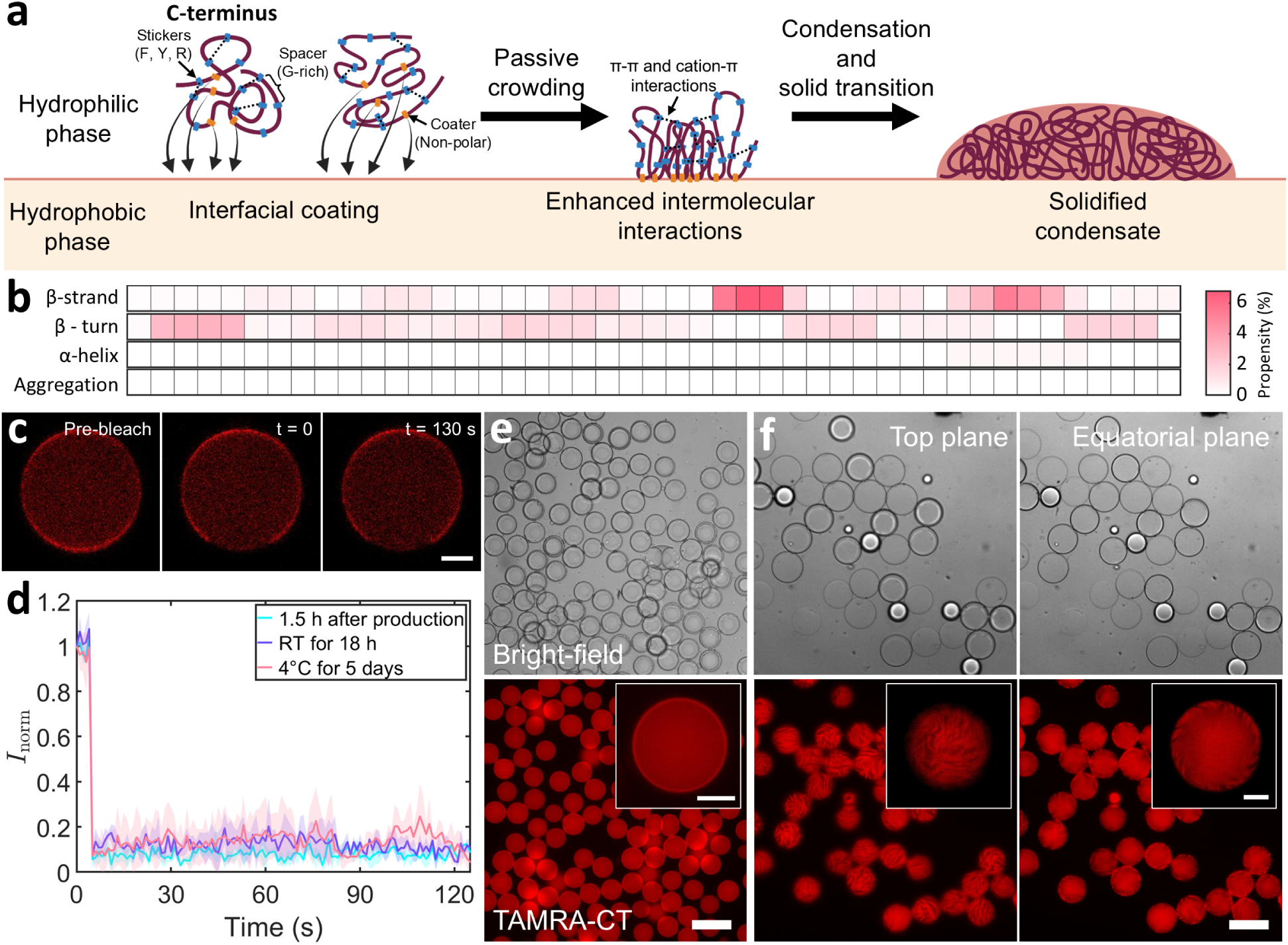
Solid transition of the CT45 molecules due to the interfacial self-assembly. **(a)** Conceptual sketch illustrating the mechanism of CT45 solid transition at the hydrophilic-hydrophobic interface. CT45 molecules are adsorbed at the interface due to the presence of the ‘coater’ amino acids. Under the accumulation, the coated CT45 molecules are passively crowded promoting the intermolecular interactions, leading to condensation, and subsequently, to the formation of a solid shell. **(b)** Secondary structure and aggregation propensity heatmap of the CT45 sequence showing the percentage of (from up to bottom) *β*-strand conformation, *β*-turn conformation, *α*-helical conformation, and aggregation. Darker colors indicate regions with higher propensity, while lighter colors represent lower propensity. **(c)** Time-lapse images taken by CLSM showing lack of fluorescence recovery of the bleached area even after 2 minutes. Scale bar, 10 *µ*m. **(d)** Plot showing lack of fluorescence recovery after bleaching under varying incubation conditions. Data points represent mean values while the shaded areas indicate standard deviations (n = 3 DEs for each condition). **(e)** Bright-field (top) and corresponding fluorescence (bottom) images showing the DEs with intact CT45 shells. **(f)** Bright-field (top) and corresponding fluorescence (bottom) images showing broken CT45 shells after stimulation by a hypotonic trigger; insets illustrate the close-ups of the broken CT45 shell with zebra-patterned cracks. For e-f, scale bar: main, 100 *µ*m; inset, 20 *µ*m. For all experiments, the encapsulated mixture consisted of 55 *µ*M CT45 (CT45:TAMRA-CT45 = 10:1, molar ratio) and 300 mM Na_2_HPO_4_, and the oil phase was fluorinated oil (HFE 7500) containing 2% FluoSurf-C surfactant. Before stimulation, DEs were suspended in the outer aqueous solution consisting of 300 mM Na_2_HPO_4_ and 1% v/v tween-20. In f, the images were captured 17 h after the hypotonic trigger (adding twice the volume of a salt-free outer aqueous to the DE suspension).

However, the rapid solidification may also be due to the *β*-sheet formation, as in such crowded environments aggregation may also be promoted. To get an indication about the aggregation potential of CT45, we used TANGO algorithm ^27^ to predict the *β*-sheet aggregation propensity of the CT45 sequences under our experimental conditions (Figure 3b): 55 *µ*M CT45 in an environment with 0.9 M ionic strength (300 mM Na_2_HPO_4_), at pH 7.4 and 25*^◦^*C. In the *β*-strand secondary structure map (Figure 3b, first panel), only the residues 26-40 exhibit limited *β*-strand propensity (lower than 6.8%), indicating overall a very low tendency for aggregation. Similarly, the probability of each residue forming *β*-turns (Figure 3b, second panel) was generally low (no higher than 3.1%), further supporting the limited aggregation potential. Moreover, the *α*-helix propensity was nearly zero across all residues (Figure 3b, third panel), consistent with the intrinsically disordered nature of the CT45 region and its high propensity for condensation. Finally, the aggregation propensity heatmap for the CT45 sequence was uniformly zero (Figure 3b, fourth panel), clearly indicating the absence of aggregation tendency. To investigate the effect of passive crowding, we increased the CT45 concentration by 100-fold (as 5.5 mM), but the aggregation propensity remained virtually to zero (Supplementary Figure 2). These findings suggest that the solid transition of CT45 at the oil–water interface is driven primarily via *π*-based interactions.

Next, we carried out FRAP experiments by confocal laser scanning microscopy (CLSM) to verify the solid nature of the CT45 structure accumulated at the interface. We started with freshly produced DEs, about 1.5 h after production, and bleached a small CT45 region at the DE interface (Figure 3c, middle). Complete absence of fluorescence recovery for over two minutes (Figure 3c, right) indicated that the CT45 shell had already transitioned to a solid-like state. Incubation of DEs at room temperature for 18 h or storing them at 4 °C for 5 days gave similar FRAP results (Figure 3d and Supplementary Figure 3), confirming that the liquid-to-solid transition of CT45 was indeed effectively completed rapidly, on the order of a few hours, without any extra trigger.

We then further probed the solid nature of the CT45 shell by expanding the DEs via a hypotonic trigger. We produced a stable, monodisperse batch of DEs displaying CT45 shells (Figure 3e, imaged after storing them for one day at 4*^◦^*C). As can be seen from the inset in Figure 3e, TAMRA-CT45 was observed to be uniformly distributed on their inner surfaces. We subsequently subjected the DEs to a hypotonic bath. Due to the osmotic pressure imbalance, water entered the DEs leading to its expansion as well that of the surrounding oil layer until equilibrium reached in a few hours (Supplementary Figure 4).^5,25^ After exposing to hypotonic bath for 17 h, while the DEs remained structurally intact, fluorescence imaging revealed the presence of zebra pattern-like multiple cracks in the CT45 shell (Figure 3f and Supplementary Movie 1). The cracks extended across the entire surface of the shell (see Supplementary Movie 2), as shown in both the top (Figure 3f left) and equatorial (Figure 3f right) planes, indicating that the CT45 molecules, being a part of a continuous shell, were unable to expand along with the extending oil surface and the shell instead cracked over multiple points to render a zebra-like pattern. This again strongly hints to the solidification of the CT45 shell adhered to the inner surface of the oil phase, rupturing as a result of the expansion. Thus, we demonstrated that the interfacial CT45 solid transition triggered by passive crowding cannot be reversed by alleviating crowded conditions, underscoring the irreversibility of this process. At the same time, the zebra-patterned cracks indicate that the CT45 shell possess multiple stress-activated weak points, exposing its fragile nature. To overcome that, we subsequently directed our efforts toward reinforcing the CT45 shell.

### Concentration-induced CT45 shell reinforcement

In order to reinforce the CT45 shell, we decided to enhance CT45 condensation throughout the vesicle lumen. To this end, we exposed DEs to a hypertonic bath to drive water out via the osmotic gradient, thereby concentrating the encapsulated contents (55 *µ*M CT45 and 300 mM Na_2_HPO_4_) and thus facilitating further condensation (see Figure 4a for a conceptual sketch). The formed condensates would then readily adhere to the interfacial CT45 shell upon contact, progressively reinforcing the shell over time. Indeed, when subjected to different osmotic pressures, the CT45 shells exhibited distinct morphological characteristics. As shown in the first panel in Figure 4b, mixing the DE suspension with an equal volume of 1 M NaCl feeding aqueous (FA, ≈2.2 times the osmotic pressure inside) led to the appearance of multiple bright spots on the CT45 shell surface (highlighted in the inset of Figure 4b), indicating that free CT45 in lumen formed condensates as a result of concentration-driven effects. With the gradual increase of the osmotic pressure (left to right in Figure 4b) up to 4 M NaCl (≈8.9 times the osmotic pressure inside), the CT45 shell surface became smoother, suggesting that the formed condensates achieved complete coverage of the initial shell. Noticeably, the forming CT45 shells also underwent significant shrinkage, indicating that the initial thin shells formed prior to stimulation were compressed and became denser (see Supplementary Movie 3). The extent of this shrinkage progressively increased with the osmotic pressure, resulting in further reinforcement of the shell. Thus, the enhancement of the CT45 shell was achieved both by depositing the internal free CT45 as well as the densification caused by the shrinkage of the shell itself.

**Figure 4:**
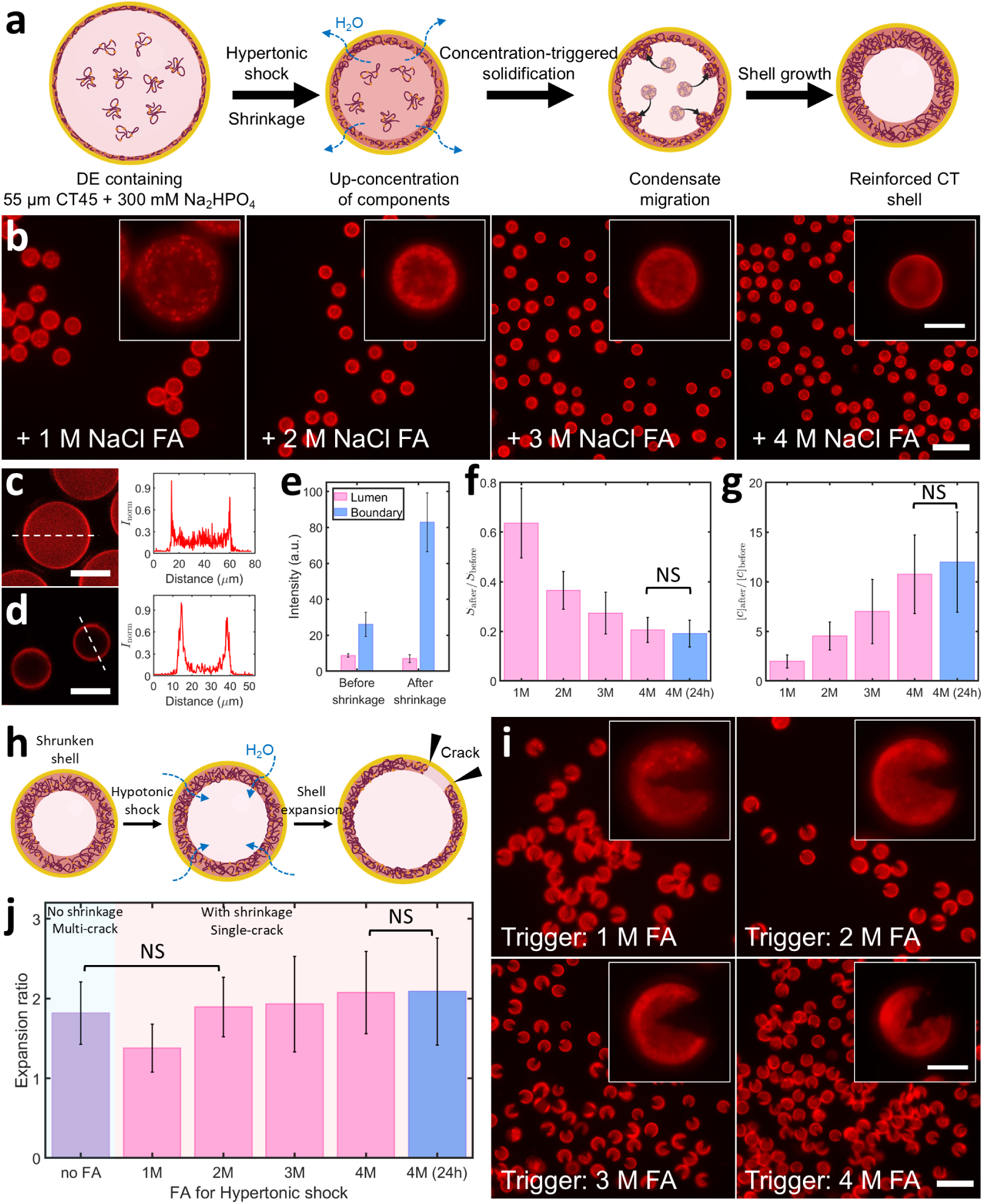
Reinforcement of the CT45 shell via hypertonic trigger. (**a**) Conceptual sketch showing DE shrinkage, enhanced CT45 condensation and subsequent shell growth. (**b**) Fluorescence images of CT45 shells after the hypertonic trigger; the insets show representative close-ups of the shell morphology after increasing the degree of the hypertonic trigger, resulting in decreasing shell sizes and smoother surfaces. Scale bar: main, 100 *µ*m; inset, 20 *µ*m. (**c-d**) Confocal images of the CT45 shell (left) and normalized fluorescence intensities (right) corresponding to the dotted lines in each panel before (c) and after (d) stimulation by 4 M FA, showing the diminished fluorescence in the lumen after shrinkage. Scale bar: 100 *µ*m (**e**) Comparison of the lumen and shell TAMRA-CT45 intensities before and after shrinkage, showing significantly higher fluorescence on the shell afterwards; *n* = 5 shells. (**f**) Normalized surface area showing the shrinkage of the CT45 shells after stimulation with different hypertonic FA. (**g**) Estimated increase in the encapsulated CT45 concentration after stimulation with different hypertonic FA, calculated based on the volume changes. For b, f, g, data were collected 1 h after the trigger, *n ≥* 70 shells. (**h**) Conceptual sketch showing the expansion of the CT45 shell after a subsequent hypotonic stimulation, leading to water influx and DE expansion, resulting in shell breakage, while keeping the DE intact. (**i**) Fluorescence images of the broken reinforced CT45 shells after the hypotonic trigger; insets illustrate representative close-ups of the broken CT45 shells, each showing a single crack. Scale bar: 100 *µ*m; inset, 20 *µ*m. (**j**) Expansion ratio of surface area showing an increase after various degrees of hypotonic triggers: pale cyan (corresponding to the CT45 shells expanded from the states in Figure 3e) and pink (corresponding to the CT45 shells expanded from the states in panel b) backgrounds respectively indicate multiple and single cracks while the horizontal axis represents the initial hypertonic triggers before the shell reinforcement. For i-j, data were collected 4 h after the trigger, *n ≥* 51 shells, NS represents not significant with p-value *>* 0.05. For all experiments, the encapsulated mixture consisted of 55 *µ*M CT45 (CT45:TAMRA-CT45 = 10:1, molar ratio) and 300 mM Na_2_HPO_4_, and the oil phase was fluorinated oil (HFE 7500) containing 2% FluoSurf-C surfactant. Before the hypertonic stimulation, DEs were suspended in the outer aqueous solution containing 300 mM Na_2_HPO_4_ and 1% v/v tween-20. In a-g, the hypertonic environment was triggered by combining the DE suspension with an equal volume of FA containing 1% v/v tween-20 and NaCl (1–4 M). In h-i, the second, hypotonic trigger was applied to the shrunken DEs obtained in b, by adding twice the volume of a salt-free outer aqueous buffer to the DE suspension. For all graphs, data points represent mean values while the error bars indicate standard deviations.

Confocal images in Figure 4c-d further demonstrate the consumption of free CT45 molecules within the lumen. Prior to the hyperosmotic shock, the interior of the CT45 shell exhibited detectable fluorescence (Figure 4c). However, after exposure to the 4 M NaCl bath, the interior fluorescence signal was largely eliminated (Figure 4d). Figure 4e compares the fluorescence intensities between the two cases. The CT45 fluorescence within the lumen diminished after the shrinkage, whereas the intensity at the bright ring boundary, representing the CT45 shell, was enhanced ≈3-fold. Meanwhile, broadening of the fluorescence peaks at the periphery supports the deposition of CT45 condensates at the shell. We further quantified the decrease in the shell surface area (Figure 4f) and the increase in the lumenal CT45 concentration (Figure 4g) by measuring the diameter of the CT45 shells before and after the triggers. Noticeably, the response of the DEs to the triggers was rapid, as no significant difference in the surface area and internal CT45 concentration (*n* ≥ 93 CT45 shells, *α* = 0.05) was observed between samples treated for 1 h (in case of 4 M NaCl trigger; pink bar in Figure 4f) and those treated for 24 h (blue bar in Figure 4g). In addition, our FRAP result also demonstrated a quick solid transition after shrinkage. As Supplementary Figure 5 shows, no fluorescence recovery was observed after 2 minutes when we bleached a region of the CT45 shell 1 h post a 4 M NaCl treatment.

Utilizing a similar DE expansion method as in Figure 3f, we subsequently evaluated the CT45 shell reinforcement. As illustrated in Figure 4h, we applied a second trigger (1 or 24 h post the first trigger indicated in Figure 4a) using a salt-free solution. Regardless of the degree of shrinkage after the first stimulation, DEs consistently expanded over the course of 4 h (Supplementary Figure 6) and stretched the confined CT45 shells until they broke to reveal a single crack, forming a Pac-Man-like structure (Figure 4i and Supplementary Movies 4-5), while keeping the DE intact (see bright-field images in Supplementary Figure 7). Compared to the zebra patterns obtained in Figure 3f, the occurrence of the single crack strongly indicates the effective reinforcement of the CT45 shell. Also, as shown in Figure 4i (corresponding to all bars with a pink background in Figure 4j), the shells consistently fractured with a single crack for all the feeding aqueous solutions. Even though the DEs treated by 2 M NaCl had the same expansion ratio 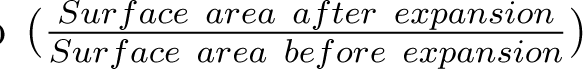 as the DEs expanded without initial shrinkage (DEs shown in Figure 3f; purple bar in Figure 4j), the shells showed distinct crack patterns. Thus, the occurrence of a single crack was independent of the degree of DE expansion. Additionally, CT45 shells that were reinforced for 1 h (pink bar) and 24 h (blue bar) both showed a single crack. In conclusion, we demonstrated concentration-induced CT45 shell reinforcement at the polar-nonpolar interface, as evidenced by an increased TAMRA-CT45 signal, thickening of the shell, and a distinct change in the fracturing pattern.

### CT45 assembly and mesoscopic scaffolds at transient interfaces

Building on the solidification and reinforcement properties of CT45 condensates at a stable oil–water interface, we explored whether a similar scaffolding effect could occur at a more transient polar–nonpolar interface. If successful, one could eventually remove the less polar organic solvent and end up with an all-aqueous system with a solid and stable proteinaceous scaffold. We began with an acetone-water system; while acetone and water are miscible solvents, owing to their polarity differences, one can still establish a transient polarity gradient before the two solvents fully mix (Figure 5a, left). This gradient can create a solvent environment that balances hydrophobic and hydrophilic interactions, thereby minimizing the overall transfer free energy of the peptide and promoting its localization.^16^ Thus, amphiphilic CT45 could preferentially accumulate in the intermediate solvent region, where the balance of water (strong hydrogen bonding) and acetone (moderate polarity) provides an optimal environment (Figure 5a, right). Indeed, upon mixing aqueous CT45 solution into acetone, we not only obtained water droplets in acetone, but immediate CT45 accumulation at the droplet interface (Figure 5b). Since acetone evaporates rapidly, when carrying out this experiment on an open glass slide, the forming CT45 shells broke within a few minutes due to the evaporative flow and a rapidly diminishing interface, leaving behind fragments (Supplementary Movie 6 and Supplementary Figure 8). We were able to form a similar shell-like CT45 accumulation by using other polar solvents, ethanol and isopropanol (see Supplementary Figure 9), as it is also less polar than water. Importantly, owing to the miscibility of acetone and water, no emulsions could be formed in the absence of CT45, underpinning the crucial role played by CT45 in stabilizing these transient interfaces. Therefore, spontaneous accumulation of CT45 at the gradient interface were critical in establishing the foundational framework that underpinned subsequent structural reinforcement.

**Figure 5:**
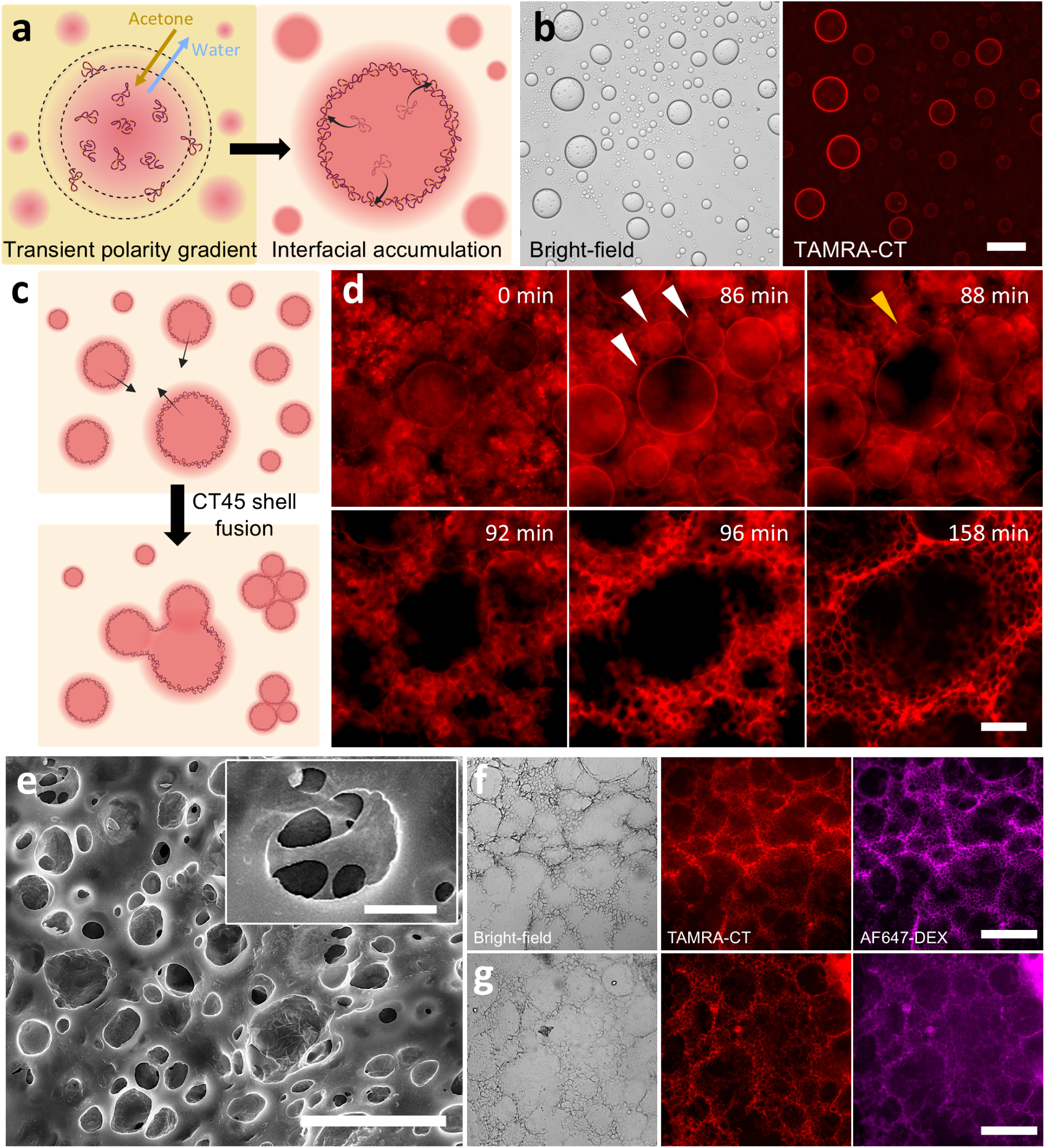
Polarity gradient-induced CT45-based scaffold formation. (**a**) Conceptual sketch showing concentration gradient induced by acetone and water, with CT45 accumulation at the interface. **(b)** Bright-field and fluorescence images showing water-inacetone droplets with immediate CT45 accumulation at the interface. Scale bar, 100 *µ*m. **(c)** Conceptual sketch showing the drying of the liquid causes the CT45 vesicles to aggregate into clusters while some CT45 shells continue to display liquid-like characteristics during this process. **(d)** Fluorescence time-lapse images showing CT45 accumulation and solidification throughout the acetone-water interface, ultimately forming a porous, scaffold-like structure. Owing to the slow evaporation of acetone, accumulated CT45 shells (pointed by the white arrows) eventually fused (pointed by the yellow arrow) and solidified to form an interconnected scaffold over the course of 3 h. Scale bar, 50 *µ*m. **(e)** Scanning electron microscopy image showing the porous surface and internal structure (inset) of the formed CT45-based scaffold structure. Scale bar: 30 *µ*m; inset, 5 *µ*m. **(f)** Bright-field and fluorescence microscopy images showing the final scaffold formed after a 3-hour drying process. The added AF647-DEX was immobilized on the CT45 scaffold. **(g)** Bright-field and fluorescence microscopy images showing that the formed structure persists after rehydration, with the dextran remaining colocalized with TAMRA-CT45, thereby confirming the structural stability of the scaffold. For f-g, scale bar, 200 *µ*m. In b, 4 *µ*L protein solution containing 55 *µ*M CT45 (CT45:TAMRA-CT45 = 10:1, molar ratio) and 300 mM Na_2_HPO_4_ was added into 36 *µ*L acetone on a clean cover glass. In d, f-g, 90 *µ*L acetone and 10 *µ*L protein solution (same composition as above with added 5 mM DEX (DEX:AF647-DEX = 1000:1, molar ratio)) were mixed together in a PVA-coated PDMS well and evaporation was reduced by putting a cover glass on top. In e, 270 *µ*L acetone and 30 *µ*L protein solution (CT45:TAMRA-CT45 = 10:1, molar ratio) were mixed together in a PDMS well with a cover slide on the bottom, and evaporation was reduced by putting a cover glass on top. In g, a 3-hour-dried structure was rehydrated by adding 100 *µ*L MQ.

Based on the observations above, the need to regulate the evaporation rate and water-acetone emulsion stability in order to maximize the liquid-to-solid transition of the interfacial CT45 became clear. To decelerate the evaporation process, we conducted the experiment in a polydimethylsiloxane (PDMS) well, overlaid with a glass slide (see Methods for details). In parallel, we added polyvinyl alcohol (PVA), an amphiphilic surfactant^28^ that may colocalize with CT45, to enhance the emulsion stability. In such a setting, the drying process of a mixture of 90 *µ*L acetone and 10 *µ*L aqueous containing 55 *µ*M CT45 (CT45:TAMRA-CT45 = 10:1, molar ratio), 300 mM Na_2_HPO_4_ and 2 % (w/v) PVA continued for ≈3.4 h and eventually formed a porous structure (Supplementary Figure 10). This indicated a positive effect of a closed environment and PVA as a surface-stabilizing agent during the CT45 scaffold formation. To further enhance its effect, we used PVA to coat the well surface in subsequent experiments (see Methods for details), rendering it hydrophilic to prevent CT45 adsorption. Meanwhile, PVA could still appear in the protein solution to aid in emulsion stabilization.

Drawing on the method established above, we further checked the ability of the formed porous structure to partition and sequester guest molecules. We added 5 mM dextran (DEX) and Alexa Fluor 647 (AF647)-labeled DEX in a molar ratio of 1000:1 as guest molecules; the latter was added for fluorescent visualization. As Figure 5d shows, the incorporation of DEX did not alter the formation of the CT45 porous structure, and the dynamic construction processes remained comparable to that observed prior to its addition. Interestingly, at the onset of the slow evaporation process, some CT45 shells continued to display liquid-like characteristics (Figure 5c). Highlighted by the arrows in Figure 5d, three CT45 shells (indicated by white arrows in frame 86 min) readily fused with each other upon physical contact, followed by their relaxation into a bigger shell structure (indicated by yellow arrows in frame 88 min). This further indicates that the fragility of CT45 shells solely induced by polarity gradient (Figure 5b) may also be due to the incomplete solidification. However, after the initial accumulation at the interface, the ongoing evaporation of acetone led to a continued increase in the concentrations of salt and CT45 in the solution. This concentration-driven condensation further facilitated solidification and reinforced the CT45 condensate shell at the interface. The decreasing CT45 fluorescence intensity inside the bright boundaries (Figure 5d) indicates the consumption of the free CT45 molecules. Consequently, after complete evaporation of acetone and water, a stable, dense and interconnected porous network was left behind (Figure 5d, frame 158 min). During the whole process, DEX fluorescence showed overlap with CT45 (see Supplementary Movie 7 and Supplementary Figure 11), indicating its clear partitioning in the CT45-rich phase. We also conducted scanning electron microscopy of the dried scaffold that clearly showed the porous structure of the formed scaffold (Figure 5e, see Methods for details). As Figure 5f shows, after 3 h of evaporation, the CT45 fluorescence was localized at the interfaces between cavities of varying sizes. This skeletal framework, along with the associated distribution of DEX, demonstrated promising structural stability even after rehydration by water, as the structure remained unaltered (Figure 5g), also proving its irreversible nature.

Overall, we demonstrated emulsion-templated CT45-based scaffolding via simple mixing of solvents with different polarities and demonstrated the capability of the scaffold to partition guest molecules.

## Discussion and Conclusion

Given its proposed role in promoting tick adhesion through LLPS, we investigated the material properties of a tick salivary protein for potential applications. We particularly focused on the key LLPS-promoting C-terminus part of the peptide (CT45), demonstrating successful simplification of the system, providing valuable insights for further bottom-up synthetic bioinspired systems. We showed the inherent nature of the CT45 peptide for interfacial accumulation and crowding, facilitated by its amphiphilic properties. This spontaneous accumulation caused CT45 to solidify under conditions that typically do not trigger phase separation, transitioning from a homogeneous to a condensed and finally to a solid state within a few hours. These findings offer new avenues in using the material potential of disordered peptides and can be broadly applied to other naturally derived proteins ^29^ or synthetic molecules.^30^

While biomolecular condensates usually behave as liquid droplets, here we leverage the ability of the CT45 peptide to undergo liquid-to-solid transition to form micrometer-scale condensate shells with aqueous cores in a controllable manner. Recent studies have explored protein-based nanovesicles^16^ by nanoprecipitation,^31^ using mixed solvents to steer precipitation-based self-assembly. In comparison, one drawback of the double emulsions templated method here could be the presence of an oil shell around individual condensate shells. However, our approach offers excellent encapsulation of biologicals and optimal control over the protein shell size in the micrometer range. Furthermore, its mechanical properties can be reinforced by controlling the shrinkage of the double emulsions. It is foreseeable that for drug delivery or cell-interaction applications, the current use of fluorinated oil could be substituted with other biocompatible alternatives, ^32^ or a water-in-oil single emulsion system could be adopted to simplify the oil phase removal.^33^ Notably, given that condensate shells have demonstrated superior delivery efficiency for biopharmaceuticals compared to condensate droplets,^17^ the presented CT45 shells hold significant promise for applications in drug delivery, food science, biomedical applications, and the construction of functional synthetic cells.

The interfacial affinity of CT45 peptide arises from the dispersed distribution of polar and nonpolar residues within its sequence. Here, we demonstrated that this interfacial affinity is not confined to well-defined stable boundaries between immiscible phases (such as water and oil), but also extends to gradient interfaces between miscible phases (such as water and acetone), where no distinct phase boundary is present. If further exploited, such peptides can be efficiently directed and be promoted to accumulate at transient interfaces, providing a potential approach to produce protein-based scaffolds, for example, useful in systems that require the removal of non-aqueous phases.^22^ By elucidating the self-assembly of an amphiphilic peptide under stable or transient polarity gradients, this study offers a mechanistic framework with significant implications for the future design and application of condensate-based vesicle systems.

In addition to the individual CT45 condensate shells, the interconnected scaffolds are also of great significance. In contrast to cytoskeleton constructed from DNA, ^34–36^ actin,^37–39^ or polymers,^40^ the CT45-based framework is remarkably minimal: the structure is constructed solely by a short peptide. This minimal polypeptide scaffold is especially valuable for streamlining the design of synthetic cells by decreasing the encapsulated contents, particularly those designed to encapsulate large amounts of internal material for diverse purposes. ^5^ Due to the irreversibility of its formation, this CT45 solid network is optimal for applications that need to respond to external stimuli and persist in a dynamic environment, e.g., providing a cytoskeleton within artificial antigen-presenting cells^41^ and keep the shape during T-cell activation. With the large-scale production of CT45 peptides or CT45-inspired molecules, this emulsion-templated condensate network also holds promise for applications in macroscopic scaffolds, such as bone tissue engineering. ^42^

In conclusion, we used a short peptide (comprised of 45 amino acids) present in a natural secretary system, and used its phase transition potential to form micron-sized shells and mechanically stable scaffolds. The proposed approach for condensate-based capsule formation is high-throughput and controllable. By clustering the interfacial condensates in a straightforward manner using evaporation as the driving force, the formed structures are stable and porous, which could provide interesting opportunities for building synthetic cell modules, for example, a cytoskeleton. Thus, we believe that this study provides valuable insights for protein design, bioengineering, and biomedical applications.

## Materials and Methods

### Materials

Sodium chloride, disodium hydrogen phosphate, dextran (MW 9-11 kDa, No. D9260), polyvinyl alcohol (PVA, average MW 30-70 kDa, P8136), 1-octanol (297887), glycerol (G2025), Pluronic™ F-68 non-ionic surfactant (24040032), ECO Tween® 20 (STS0200) were purchased from Sigma-Aldrich. Labeled dextran (Alexa Fluor™ 647, 10,000 MW, D22914) was purchased from Fischer Scientific B.V. Phospholipids including DOPC (SKU 850375C), DOPS (SKU 840035P) and cy5-DOPE (SKU 810335C) were purchased from Avanti Polar Lipids, Inc. Sylgard™ 184 silicone elastomer (PDMS) and curing agent were purchased from Dow. Silicon wafer was bought from Silicon Materials. Photoresist (EpoCore 10) and photoresist developer (mr-Dev 600) were purchased from Micro resist technology GmbH. Microfluidic accessories including liquid flows tygon tubing coil 1/16” OD X 0.02” ID (SKU: LVF-KTU-13), stainless steel 90° Bent PDMS couplers (SKU: PN-BEN-23G), rapid-core microfluidic punches (D = 0.5 mm, 0.75 mm and 3 mm), PTFE Tubing 1/16” OD (SKU: BL-PTFE-1608-50), and HFE 7500 fluorinated oil containing 2% FluoSurf-C surfactant (SKU: EU-FSC-V10-2%-HFE7500) were purchased from Darwin Microfluidics. Elveflow pressure controller OB1-MK3 was used to control the fluid flow.

### Peptide synthesis and fluorescent labelling

All peptides (CT45 and TAMRA-CT45) were synthesized via solid-phase peptide synthesis (SPPS), using Boc-based SPPS and Fmoc-based SPPS. Comprehensive details of the synthesis, labeling, and characterization are described elsewhere. ^11^ For the labeling of CT45, tetramethyl rhodamine-5-maleimide (Invitrogen, Molecular Probes, Eugene, Oregon, USA) was used. Details of characterization are provided in Supplementary Figures 12-13.

### Microfabrication and surface functionalization

The master wafers for droplet production were prepared by standard photolithography, ^43–45^ with the channel height kept at 10 *µ*m for liposome production^5,22–24^ and 20 *µ*m for DE production.^5,25^ The master wafer for lipid bilayer experiment was prepared by dual-layer photolithography, ^46^ to produce multi-height patterns. The first layer of the microchannel was made with a height of 20 *µ*m, while the second layer for the microwells was 30 *µ*m high. Consequently, the final structure consisted of a 20 *µ*m-high channel with multiple 10 *µ*m-high microwell patterns on top.

Based on the wafer with varied patterns, we fabricated the microfluidic chips through standard soft lithography. By pouring the PDMS mixture (10:1 weight ratio for PDMS and the curing agent) on the wafer and degassing using a vacuum desiccator, we produced the PDMS blocks with the designed channel. Meanwhile, we produced PDMS-coated glass slips by spin coating PDMS mixture on a coverglass (Corning® no. 1) at 500 rpm for 15 s (at an increment of 100 rpm/s) and then at 1000 rpm for 30 s (at an increment of 500 rpm/s). Both PDMS-covered wafer and PDMS-coated glass were baked at 70 °C for 2 h. After curing, we carefully removed the hardened PDMS blocks from the wafer and punched inlet and outlet holes using biopsy punches. For liposome production, inlet and outlet diameter was 0.5 and 3 mm, respectively; for DE production and lipid bilayer experiment, all punched holes were in diameter 0.75 mm. For liposome or DE production, we bonded PDMS block on the coverslip by treating them at 12 MHz (RF mode high) for 30 seconds using a plasma cleaner (Harrick Plasma PDC-32G). The bonded device was then baked at 70 °C for 2 h. After baking, we treated the production chips by flowing PVA solution (5% w/v, molecular weight 30 - 70 kDa) in outer aqueous channels for 15 minutes. During the treatment, we kept positive pressures on the inner aqueous and oil inlets to retain the PVA-air boundary stable at the production junction. Afterwards, we moved out all PVA solution by applying 2-bar pressure on the inner aqueous and oil inlets and a pressure of -1 bar at the exit. The devices were eventually dried on a hotplate with 120 °C for 15 minutes. For the chips of the lipid bilayer experiment, the PDMS blocks and PDMS-coated slides were saved until plasma treated before vesicle loading.

The PDMS wells for FRAP, DE treatments, and scaffold formation were prepared by soft lithography on a clean silicon wafer. We used the same approach to produce PDMS blocks by curing PDMS mixture (10:1 weight ratio with curing agent) on a wafer without any pattern. Then we punched 5 mm-holes as the wells on PDMS blocks and plasma bonded them on a PDMS-coated glass slide. The surface-functionalization of the wells was conducted after one-hour baking at 70 °C, followed by PVA treatment for 15 minutes. In the end, the devices were dried on a hotplate with 120 °C for 15 minutes.

### Lipid bilayer interaction in microchannels

We formed the nanovesicles by extrusion and spread them on the microchannel to form a lipid bilayer (see Supplementary Figure 1a). Lipids in chloroform (89.9 % DOPC + 10 % DOPS + 0.1% cy5-DOPE, molar ratio) were placed into a glass vial and then dried under a gentle nitrogen stream. The vial was then desiccated in vacuo overnight (at least 12h) to remove extra chloroform. The resulting lipid film was then rehydrated in PBS to reach a final lipid stock concentration of 1 mM. After fully rehydrating the lipid film, so that no lipid film was seen stuck to the glass vials, the hydrated lipid suspension was extruded 21 times through a polycarbonate membrane with pore sizes of 100 nm with the Avanti Polar Lipids Mini extruder. The small unilamellar vesicles (SUVs) were diluted with PBS to a final lipid concentration of 0.2 mM. Dynamic light scattering was performed to confirm the SUV size of ≈130 nm (see Supplementary Figure 14). Vesicles were stored at 4°C and used for up to one week. When making the lipid bilayer in microfluidic device, we loaded the vesicle suspension (0.2 mM total lipid concentration) into the chip 30 minutes post plasma bonding and baking. After 1 h incubation at room temperature in the dark room, we slowly flowed 10 times-concentrated PBS by applying 100 mbar on inlet for 5 minutes and washed the channel with PBS for 5 minutes with the same pressure.

CT was loaded into microfluidics via the XXS microfluidic reservoir. After the lipid bilayer formation on the microchannel surface, 10 *µ*L of CT45 solution (125 *µ*M CT45, CT45:TAMRA-CT45 = 10:1, molar ratio, in PBS) was added to the reservoir connected to the microfluidic pump. The pressure was slowly increased until liquid was seen coming out from the microfluidic outlet. After the one-hour CT45 incubation, we replaced the XXS microfluidic reservoir with a tubing connected to PBS and slowly flushed the channel using the same pressure to check the interaction of CT45 and coated lipid bilayer.

### Liposome and DE production

Both Liposomes and DEs were produced by the microfluidic platform. We used octanol-assisted liposome assembly (OLA) method^22^ to produce liposomes. Four solutions were prepared: inner aqueous (IA), outer aqueous (OA), lipids in 1-octanol (LO), and exit well aqueous (EA). IA, OA, and EA contained 300 mM Na_2_HPO_4_ and 15% v/v glycerol. Additionally, 5% w/v F68 surfactant was present in OA. Components in IA included 55 *µ*M CT45 (CT45:TAMRA-CT45 = 10:1, molar ratio). Components in LA included 89.9 % DOPC, 10% DOPS and 0.1% cy5-DOPE (molar ratio) with a final DOPC concentration of 2 mg/ml. A detailed production and collection protocol is described elsewhere. ^24^ For double emulsion production, we used a two-junction design as described previously.^25^ IA contained 55 *µ*M CT45 (CT45:TAMRA-CT45 = 10:1, molar ratio) and 300 mM Na_2_HPO_4_. OA contained 300 mM Na_2_HPO_4_ and 1% v/v Tween-20. The oil phase was HFE 7500 fluorinated oil (containing 2% FluoSurf-C surfactant). We collected produced DEs from a clean pipette tip (200 *µ*L) inserted at the outlet and stored them in a dark glass vial at 4 °C.

### Fluorescence recovery after photobleaching

For FRAP experiments, we pipetted 90 *µ*L of OA and 10 *µ*L of DE expansion into a PDMS well. The well was sealed with a cover glass to avoid evaporation. For bleaching, the regions of interest (ROI) were a square area with a side length of approximately 20 *µ*m and they were bleached using 100% laser intensity for 20 frames with a frame interval of 443 ms. The intensity of the bleached area was normalized by 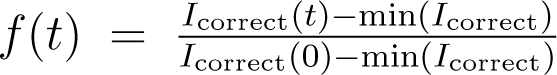, where 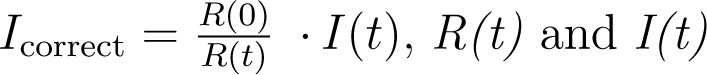 indicate the fluorescence intensity of the reference droplet at time *t* and the original fluorescence intensity of the bleached region at time *t*, respectively; min(*I_correct_)* indicates the minimum value of *I_correct_*, which is obtained right after the sample is bleached.^5^

### DE treatment and observation

For all DE experiments, we first diluted the emulsions by pipetting 35 *µ*L of OA and 5 *µ*L of the DE suspension in the PDMS well. For the experiment related to Figure 3f, we added 80 *µ*L of FA (1% v/v Tween-20 in Milli-Q) in 40 *µ*L of DE suspension in PDMS well. For the experiments related to Figure 4, we first pipetted 40 *µ*L of first FA (1% v/v Tween-20 and 1–4 M NaCl) into 40 *µ*L of DE suspension (for Figure 4b). After 1 h-incubation to allow for the shrinkage, we removed 40 *µ*L of the supernatant, and immediately refilled the PDMS well with 80 *µ*L of second FA (1% v/v tween-20 in Milli-Q)(for Figure 4i).

### Evaporation-induced CT45 scaffold

For emulsion-templated scaffold formation (Supplementary Figure 10), we made the emulsions in 5 mm-PDMS wells by pipetting 10 *µ*L of aqueous solution (55 *µ*M CT45, CT45:TAMRA-CT45 = 10:1, molar ratio, 300 mM Na_2_HPO_4_ and 2% w/v PVA, molecular weight 30 -70 kDa) into 90 *µ*L of acetone. The solutions were mixed once by pipetting up and down to avoid disrupting the scaffold formation. The well was covered by a cover slide to reduce evaporation.

For the formation of scaffold requestering guest molecules (Figure 5d), we made the emulsions in 5 mm-diameter PVA-coated PDMS wells by pipetting 10 *µ*L of aqueous solution (55 *µ*M CT45, CT45:TAMRA-CT45 = 10:1, molar ratio and 300 mM Na_2_HPO_4_) into 90 *µ*L of acetone. The solutions were mixed once by pipetting up and down to avoid disrupting the scaffold formation. The well was covered by a cover slide to reduce evaporation.

For the samples of scanning electron microscope, we used a 15mm-PDMS well and put a 12 mm round coverslip coated with poly-L-Lysine on the bottom for scaffold formation. The experiments were conducted by pipetting 30 *µ*L of aqueous solution (55 *µ*M CT45, CT45:TAMRA-CT45 = 10:1, molar ratio, 300 mM Na_2_HPO_4_ into 270 *µ*L of acetone. After one day of evaporation at room temperature, we took the coverslip with CT45 scaffold for further imaging.

### Microscopy

Images for double emulsion experiments were acquired using a ZEISS microscope (Axio Observer 7) equipped with Light Source Colibri 5 (Type RGB-UV), a ZEISS Plan-NEOFLUAR 20 x /0.5 objective, and a 90 HE LED filter set. For bright-field visualization, images were acquired at an exposure of 100 ms with 2 V light intensity while for fluorescence visualization, samples were excited with 50% light intensity and images were acquired at exposure of 100 ms, using a Prime BSI Express sCMOS camera.

Confocal and FRAP experiments were performed on Leica SP8-SMD microscope using 63x (Numerical Aperture 1.2) water objective. For bleaching, the ROI was bleached using 100% laser intensity of a tunable white light laser source (LEICA TCS SP5 X). The recovery of the bleached area was recorded for approximately 2 minutes.

### Scanning electron microscopy

The CT45 scaffold sample on coverslips was observed by scanning electron microscopy (SEM, Magellan 400, FEI, USA). For imaging, the coverslips were placed on the conductive carbon tape, and the sample surfaces were coated with a thin, fine-grained tungsten layer using a high-vacuum sputter coater (SCD 500, Leica, Germany) to enable high-resolution analysis. To facilitate observation of the internal porous structure, the holes on the surface were specifically identified and imaged.

### Image analysis

ImageJ was used for image processing and analysis in the case of FRAP experiments, fluorescence intensity analysis as well as size measurements. In case of DE experiments, only those DEs that were non-clustered and in focus were taken into account for the analysis (MATLAB R2019b). Error bars in the graphs indicate the standard deviation of the mean for respective samples.

## Data Availability

Data supporting the findings of this study are available within the paper, its Supplementary Information, Supplementary Movies 1-7, and Source Data 1. Any additional supporting data is available from the corresponding author upon reasonable request.

## Statistics and reproducibility

The double emulsion production (Figure 3e), CT45 coating (Figure 2i-j), FRAP and confocal imaging (Figure 3c-d, 4c-d, Supplementary Figure 1b-c, 3, 5), emulsion-template scaffold formation (Figure 5b, 5d Supplementary Figure 8, 10, 11) and scaffold rehydration (Figure 5f-g) experiments were repeated at least three times. The DE treatments (Figure 3f, 4, Supplementary Figure 4, 6, 7) were repeated twice for each of the five conditions. The lipid bilayer (Figure 2d) and liposome (Figure 2f-g) experiments, scanning electron microscopy (Figure 5e), ethanol/isopropanol experiment (Supplementary Figure 9) and dynamic light scattering (Supplementary Figure 14) were repeated once.

## Acknowledgments

S.D. and C.C. acknowledge financial support from Dutch Research Council (grant number: OCENW. KLEIN. 465). Schematics were created using BioRender.com.

## Author contributions

Conceptualization: C.C. and S.D.; Investigations and methodology: C.C. and S.D.; Microfluidics and microscopy: C.C.; Lipid bilayer experiment: N.J.; Scanning electron microscopy: Q.W.; Protein synthesis and labelling: I.D.; Data curation: C.C.; Formal analysis: C.C., I.D., and S.D.; Funding acquisition and project administration: S.D.; Resources: I.D., S.D.; Supervision and validation: S.D.; Visualization: C.C., S.D.; Writing (original draft): C.C, S.D.; Writing (review and editing): C.C., Q.W., N.J., I.D., and S.D. All authors have read and given their consent to the final version of the publication.

## Declarations

Authors declare no competing interests.

