## Supplementary information for "Condensate-based shells and scaffolds via interfacial liquid-to-solid transition of disordered peptides"

### Supplementary Figures

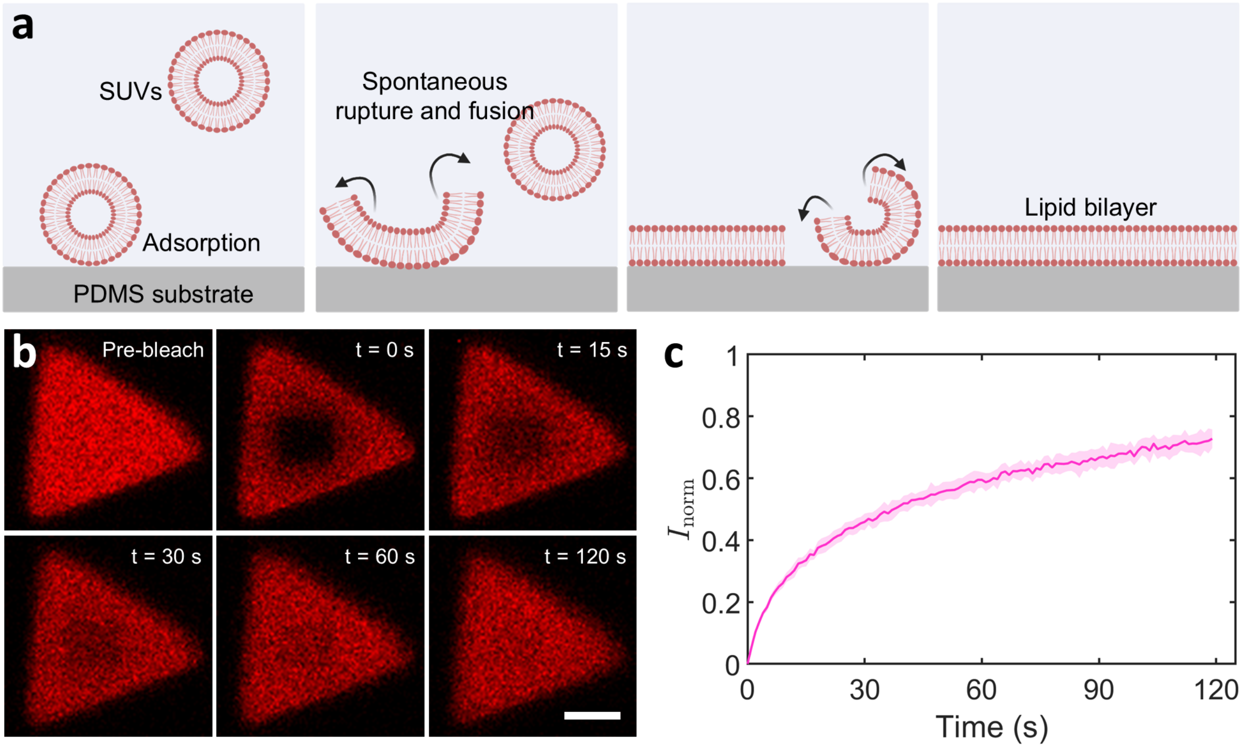


Supplementary Figure 1: **Demonstration of the supported lipid bilayer formation via FRAP experiments. (a)** Schematic of the vesicle fusion method for bilayer formation. SUVs are incubated in the plasma-treated microwells. Vesicles adsorb to the surface, rupture, fuse to the surface and self-assemble, forming a homogenous lipid bilayer. **(b)** Time-lapse CLSM images showing the FRAP experiment conducted on a triangular microchamber present in the microchannel. Scale bar, 10 $\mu$m. **(c)** Fluorescence recovery curve showing the recovery of the lipid bilayer over 2 minutes. Data points represent mean values while the shaded areas indicate standard deviations (n = 10). The lipid composition was 99.9 % DOPC + 0.1% cy5-DOPE (molar ratio) and the diffusion coefficient obtained from fitting the data ($2.8 \pm 0.3\,\mu m^2/s$, $R^2 = 0.97$) was consistent with the reported range for a 100% DOPC bilayer (2–3 $\mu m^2/s$).[1–3]

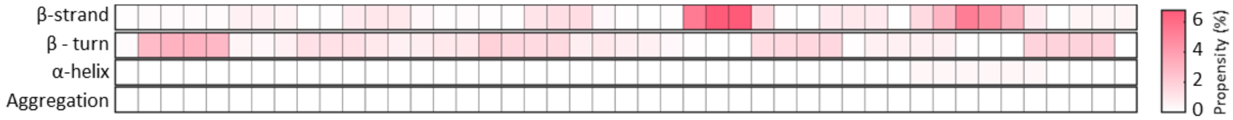


Supplementary Figure 2: **Secondary structure and aggregation propensity heatmap of the CT45 sequence with 100 times higher concentration.** From top to down: the percentage of $\beta$-strand conformation, $\beta$-turn conformation, $\alpha$-helical conformation and aggregation. Darker colors indicate regions with higher propensity, while lighter colors represent lower propensity. TANGO algorithm[4] was applied for the prediction with the conditions of 5500 $\mu$M CT45 in an environment with 0.9 M ionic strength (300 mM $Na_2HPO_4$), at pH 7.4, and 25°C.

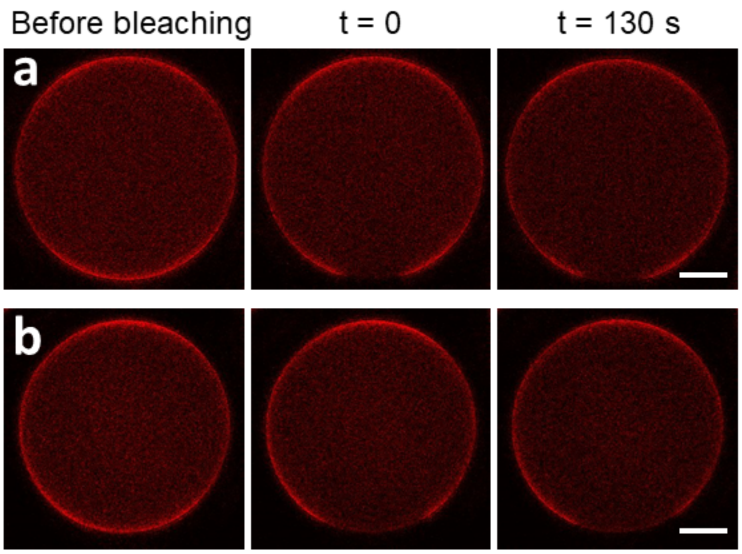


Supplementary Figure 3: **Verification of CT45 shell solidification via FRAP assay under varying DE incubation conditions.** Time-lapse CLSM images showing no fluorescence recovery of the CT45 shell for both **(a)** DEs saved in 4°C for 5 days and **(b)** DEs exposed at room temperature for 18 h. Scale bar, 10 $\mu$m. The encapsulated mixture consisted of 55 $\mu$M CT45 (CT45:TAMRA-CT45 = 10:1, molar ratio) and 300 mM $Na_2HPO_4$, and the oil phase was fluorinated oil (HFE 7500) containing 2% FluoSurf-C surfactant.

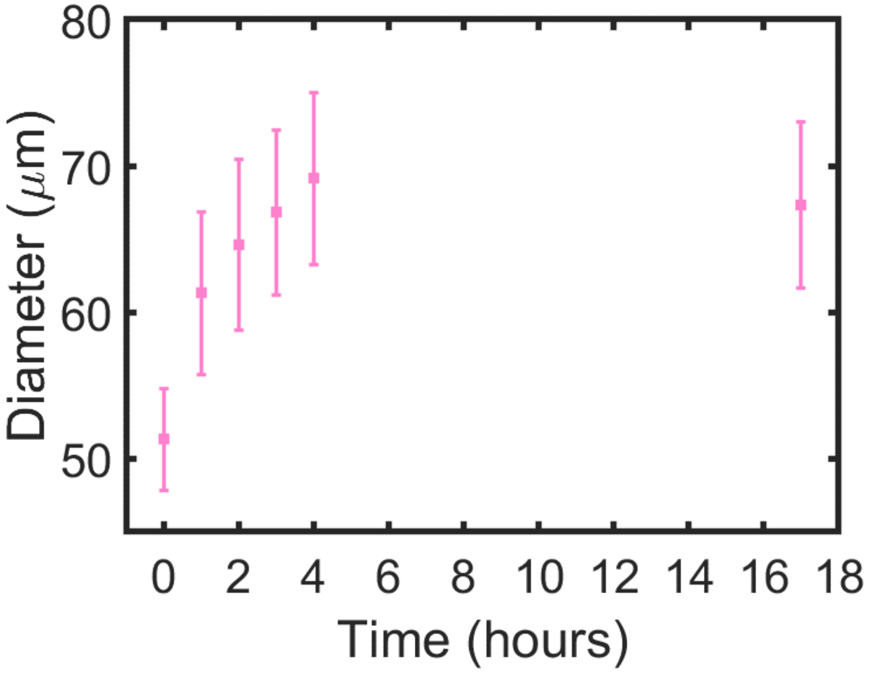


Supplementary Figure 4: **The size of the double emulsions initially expanded upon hypotonic trigger and then remained stable until 17 hours.** The size of the DEs increased from 50 $\mu$m till 70 $\mu$m within 4 hours and then remained constant. Data points represent mean values derived from $\geq$ 51 DEs, with error bars indicating standard deviations. The encapsulated mixture consisted of 55 $\mu$M CT45 (CT45:TAMRA-CT45 = 10:1, molar ratio) and 300 mM $Na_2HPO_4$, and the oil phase was fluorinated oil (HFE 7500) containing 2% FluoSurf-C surfactant. Before the hypotonic stimulation, the DEs were suspended in the OA consisting of 300 mM $Na_2HPO_4$ and 1% v/v tween-20. The hypotonic condition was triggered by combining the DE suspension with a twice volume of FA containing 1% tween-20 without NaCl.

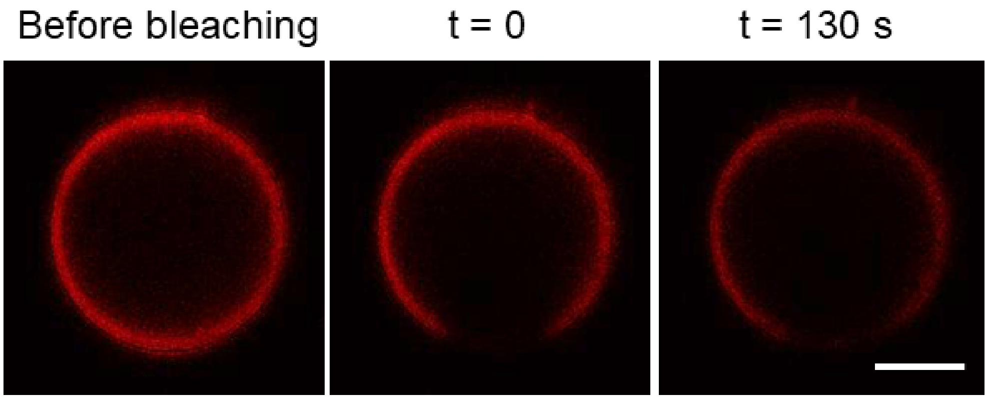


Supplementary Figure 5: **Verification of CT45 shell solidification via FRAP assay after DE shrinkage.** Time-lapse CLSM images showing no fluorescence recovery of the CT45 shell after the DEs were exposed to a 4 M-NaCl bath for 1 hour. Scale bar, 10 $\mu$m. The encapsulated mixture consisted of 55 $\mu$M CT45 (CT45:TAMRA-CT45 = 10:1, molar ratio) and 300 mM $Na_2HPO_4$, and the oil phase was fluorinated oil (HFE 7500) containing 2% FluoSurf-C surfactant. DEs were treated by combining the suspension with an equal volume of FA containing 1% v/v tween-20 and 4M NaCl.

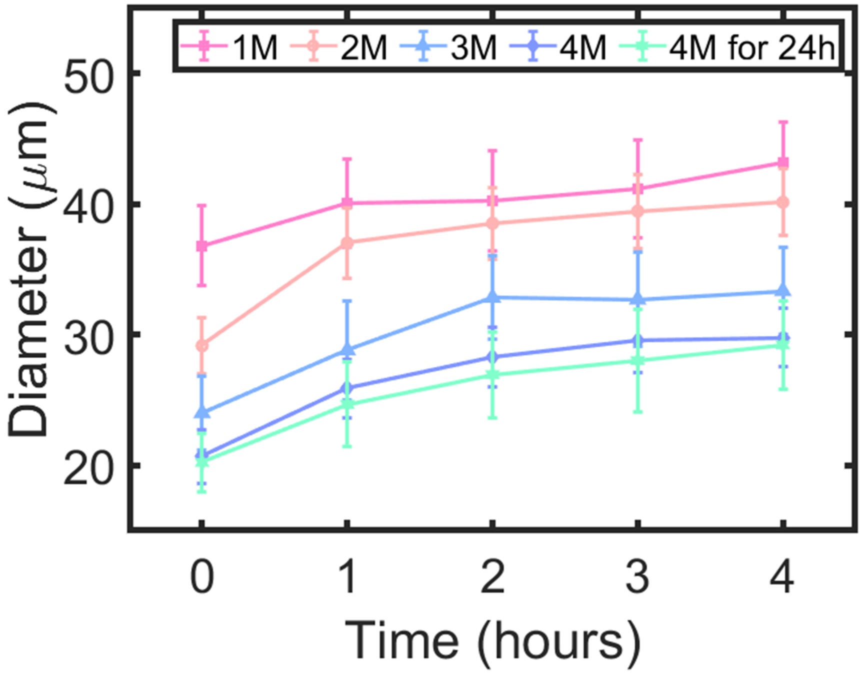


Supplementary Figure 6: **The size of shrunk DEs expanded within 4 hours after the second (hypotonic) trigger.** For various kinds of hypotonic triggers (1–4 M NaCl), the DE diameter increased for the first few hours before reaching a plateau. Data points represent mean values derived from $\geq$ 67 DEs, with error bars indicating standard deviations. For all experiments, the encapsulated mixture consisted of 55 $\mu$M CT45 (CT45:TAMRA-CT45 = 10:1, molar ratio) and 300 mM $Na_2HPO_4$, and the oil phase was fluorinated oil (HFE 7500) containing 2% FluoSurf-C surfactant. Before the first (hypertonic) stimulation, the DEs were suspended in OA solution consisting of 300 mM $Na_2HPO_4$ and 1% v/v tween-20. The first (hypertonic) trigger consisted of combining the DE suspension with an equal volume of FA containing 1% v/v tween-20 and NaCl (1–4 M). After 1 hour of incubation (24 hours for the last case), the suspensions were combined with twice the volume of a salt-free FA solution containing 1% v/v tween-20 as the second (hypotonic) trigger. The legend indicates the concentration and incubation time used for the first stimulation. If the time is not indicated, the incubation time was 1 hour.

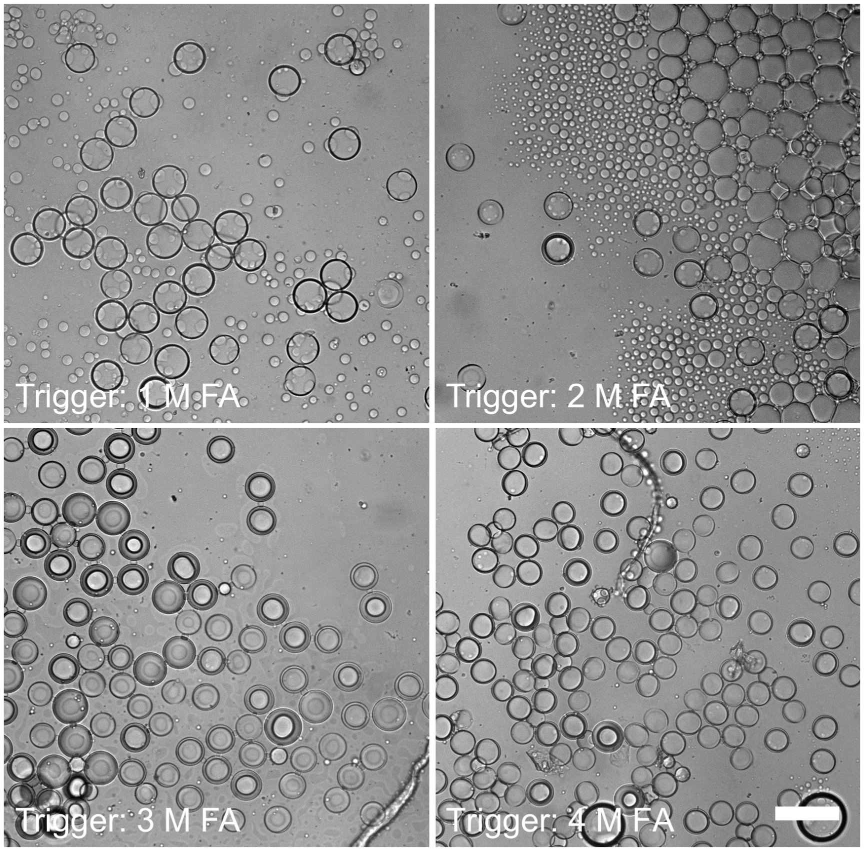


Supplementary Figure 7: **Intact DE structure after the hypotonic trigger.** Bright-field images of the intact DEs containing broken CT45 shells imaged at the equatorial plane after various hypotonic triggers. These images correspond to the CT45 fluorescence images shown in Figure 4i. The encapsulated mixture consisted of 55 $\mu$M CT45 (CT45:TAMRA-CT45 = 10:1, molar ratio) and 300 mM $Na_2HPO_4$, and the oil phase was fluorinated oil (HFE 7500) containing 2% FluoSurf-C surfactant. Before the hypertonic stimulation, DEs were suspended in OA containing 300 mM $Na_2HPO_4$ and 1% v/v tween-20. Then the hypertonic environment was triggered by combining the DE suspension with an equal volume of FA containing 1% v/v tween-20 and NaCl (1–4 M). Afterwards, the second, hypotonic trigger was applied to the shrunken DEs obtained, by adding twice the volume of a salt-free OA to the DE suspension. The images were collected 4 hours after the second trigger. The smaller droplets seen in some of the images are unwanted oil droplets, primarily formed during the pinching-off process as a result of unstable production.

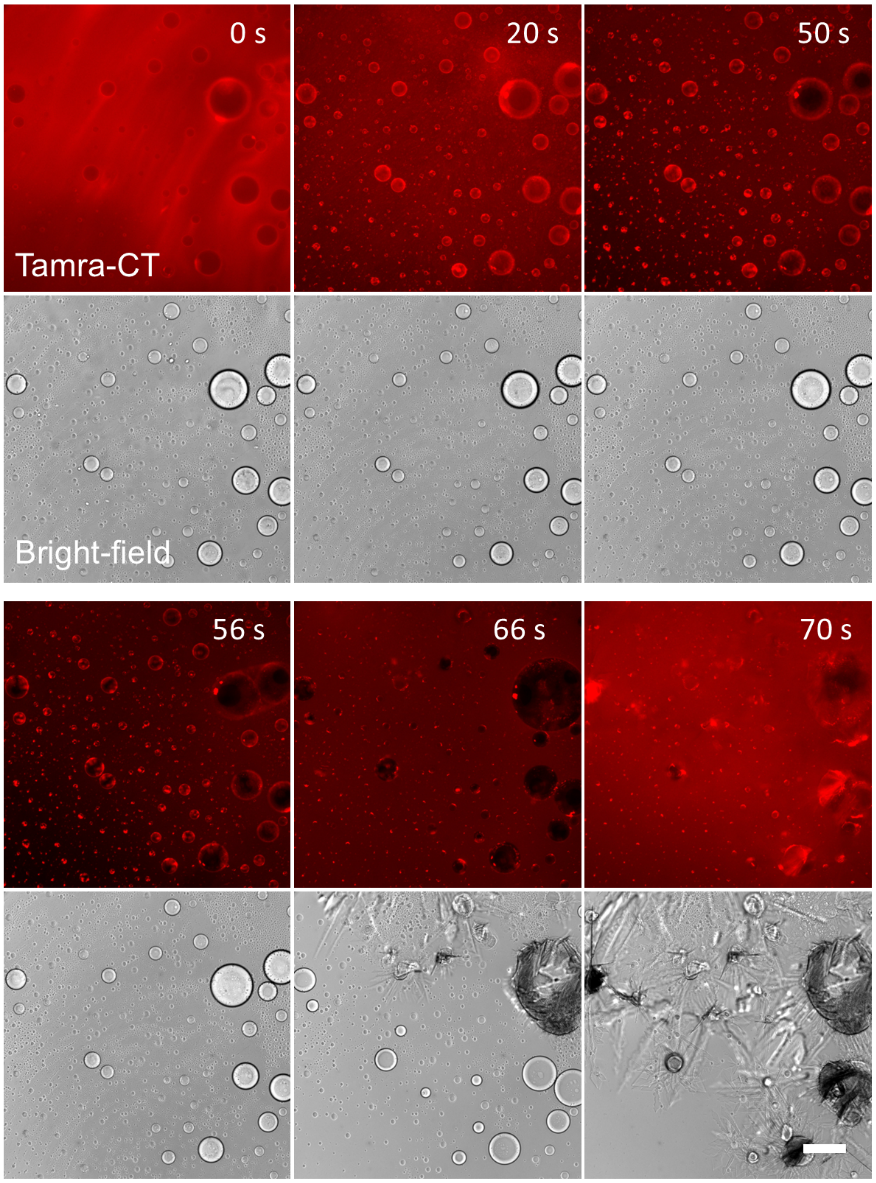


Supplementary Figure 8: **Fluorescence time-lapse images showing CT45 accumulation at the interface, shell formation and breakage after mixing acetone and CT45 solution on a glass slide.** Due to the rapid evaporation of acetone, CT45 shells were unable to form and get reinforced upon mixing and easily broke upon evaporation of acetone. Scale bar, 200 $\mu$m. The experiment was conducted by mixing 2 $\mu$L protein solution containing 55 $\mu$M CT45 (CT45:TAMRA-CT45 = 10:1, molar ratio) and 300 mM $Na_2HPO_4$ into 18 $\mu$L acetone on a clean cover glass.

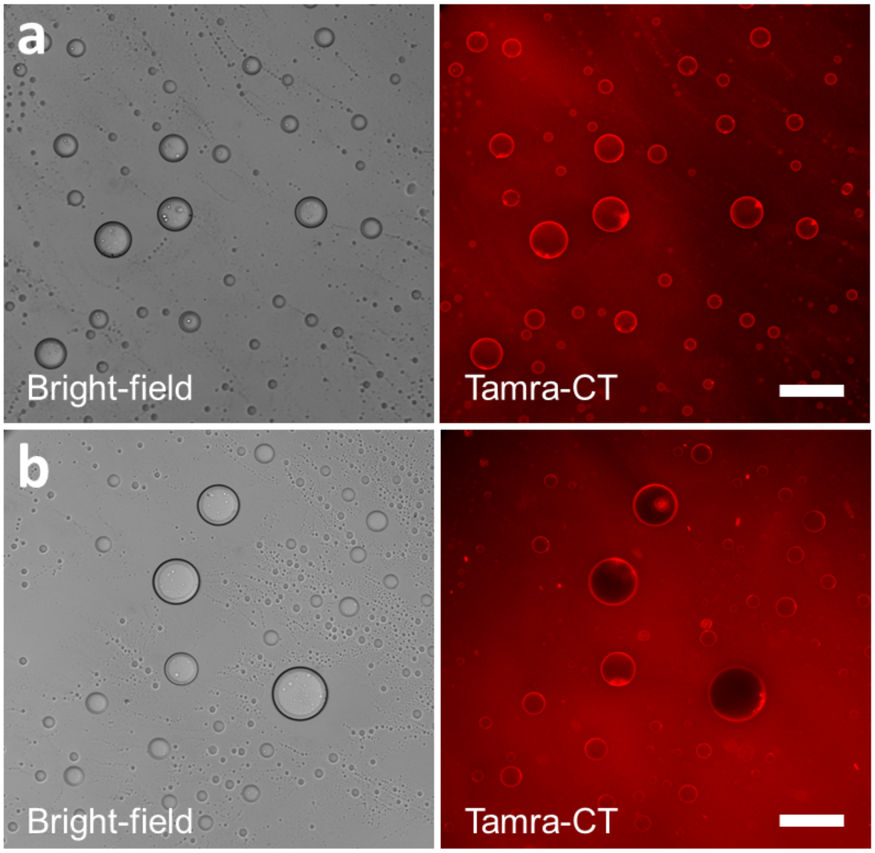


Supplementary Figure 9: **Bright-field and fluorescence images showing the self-assembly of CT45 peptides by polarity gradient using different solvents.** (a) Interfacial CT45 accumulation in presence of ethanol. (b) Interfacial CT45 accumulation in presence of isopropanol. Scale bar, 200 $\mu$m. The experiment was conducted by pipetting 2 $\mu$L protein solution containing 55 $\mu$M CT45 (CT45:TAMRA-CT45 = 10:1, molar ratio) and 300 mM $Na_2HPO_4$ into 18 $\mu$L ethanol/isopropanol on a clean cover glass.

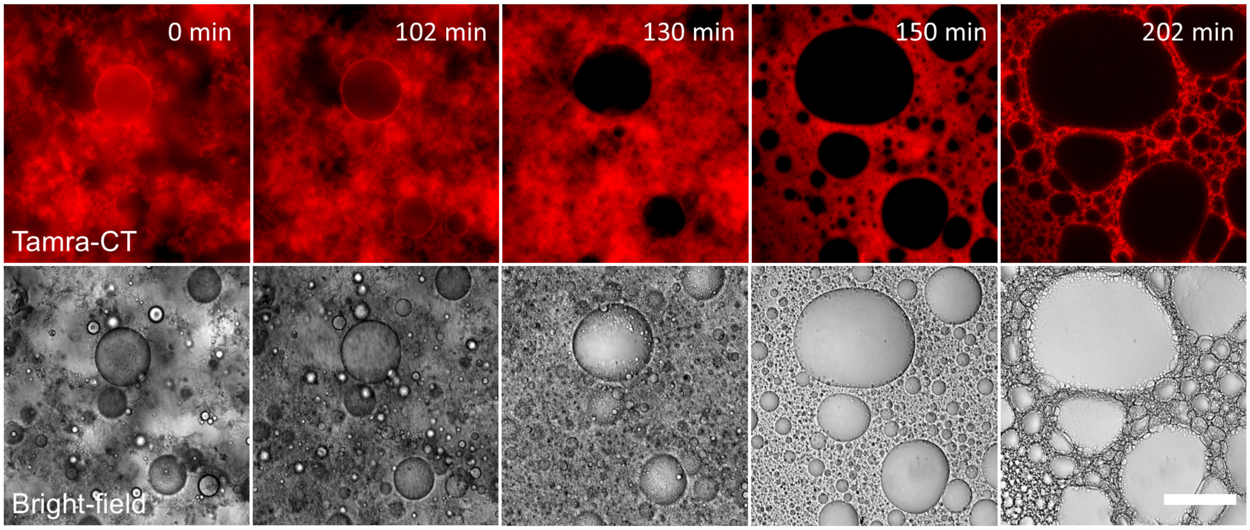


Supplementary Figure 10: **Fluorescence and bright-field time-lapse images showing the scaffold formation after mixing acetone and CT45 solution in a sealed PDMS well.** By reducing the acetone evaporation rate, CT45 gradually accumulated, solidified and clustered at the interface and eventually stabilized in the form of an interconnected structure after 3 hours. Scale bar, 200 $\mu$m. The experiment was conducted by mixing 90 $\mu$L acetone and 10 $\mu$L aqueous solution containing 55 $\mu$M CT45 (CT45:TAMRA-CT45 = 10:1, molar ratio), 300 mM $Na_2HPO_4$ and 2 % (w/v) PVA. Evaporation was significantly reduced by putting a cover glass on the clean PDMS well.

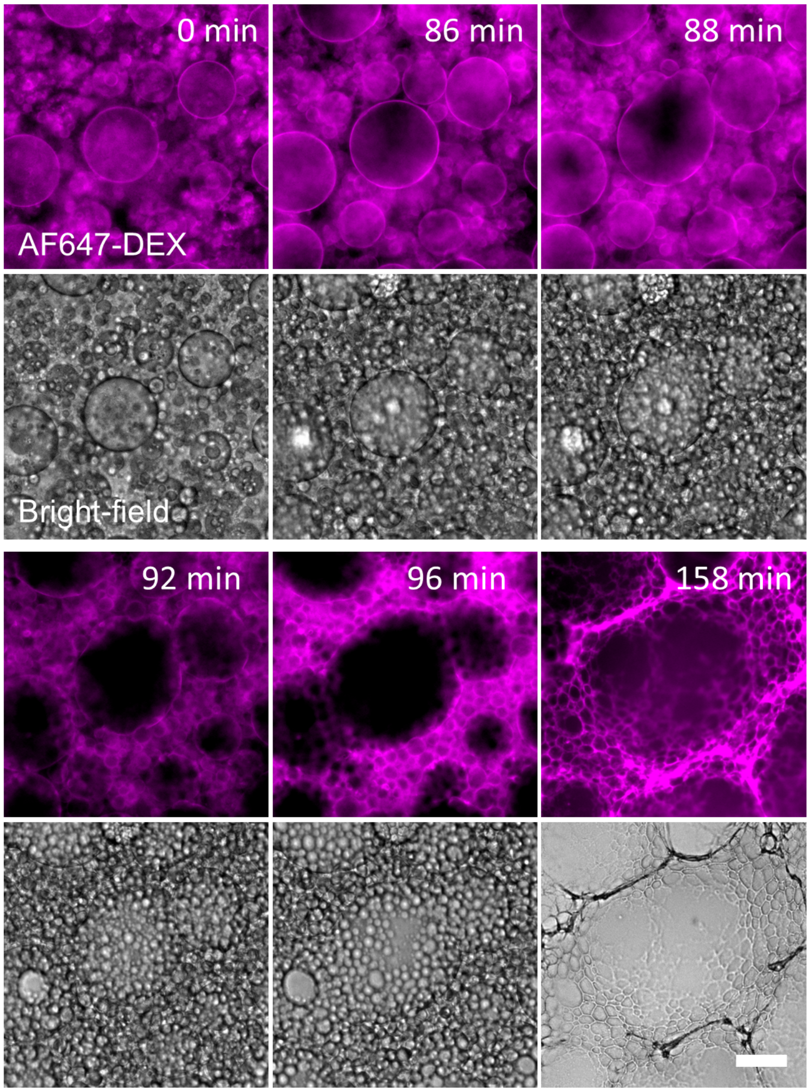


Supplementary Figure 11: **DEX effectively partitions into the CT45 scaffold.** DEX fluorescence and bright-field time-lapse images showing DEX partitioning in CT45 skeleton-like structure during its formation. The time-lapse corresponds to the CT45 fluorescence time-lapse images shown in Figure 5d. Owing to the slow evaporation of acetone, accumulated CT45 eventually solidified and formed an interconnected network over the course of 3 hours, with the DEX fluorescence overlapping with CT45 fluorescence showing its strong partitioning into the structure. Scale bar, 50 $\mu$m. To form the CT45 scaffold, 90 $\mu$L acetone and 10 $\mu$L aqueous solution containing 55 $\mu$M CT45 (CT45:TAMRA-CT45 = 10:1, molar ratio), 300 mM $Na_2HPO_4$ and 5 mM DEX (DEX:AF647-DEX = 1000:1, molar ratio)) were mixed together in a PVA-coated PDMS well and evaporation was significantly reduced by putting a cover glass on top.

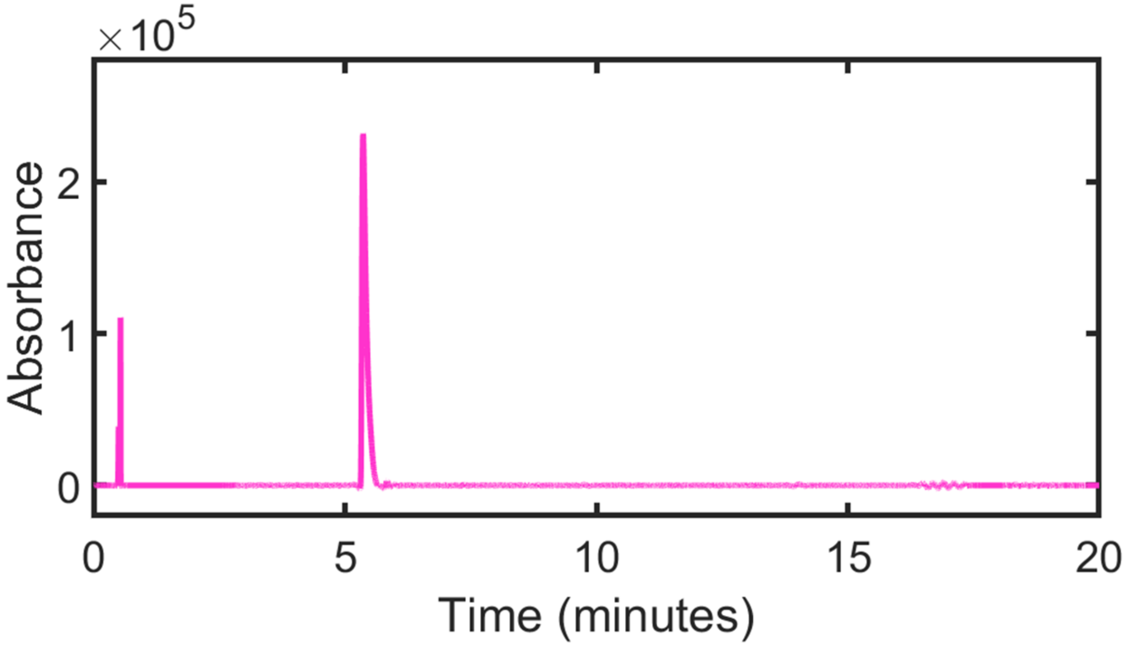


Supplementary Figure 12: **HPLC chromatogram of TAMRA-CT45.** HPLC trace of the purified TAMRA-CT45 synthesized using Fmoc-based SPPS (220 nm). The injection peak is visible at 0.04 minutes and the retention time of the protein is approximately 5.4 minutes.

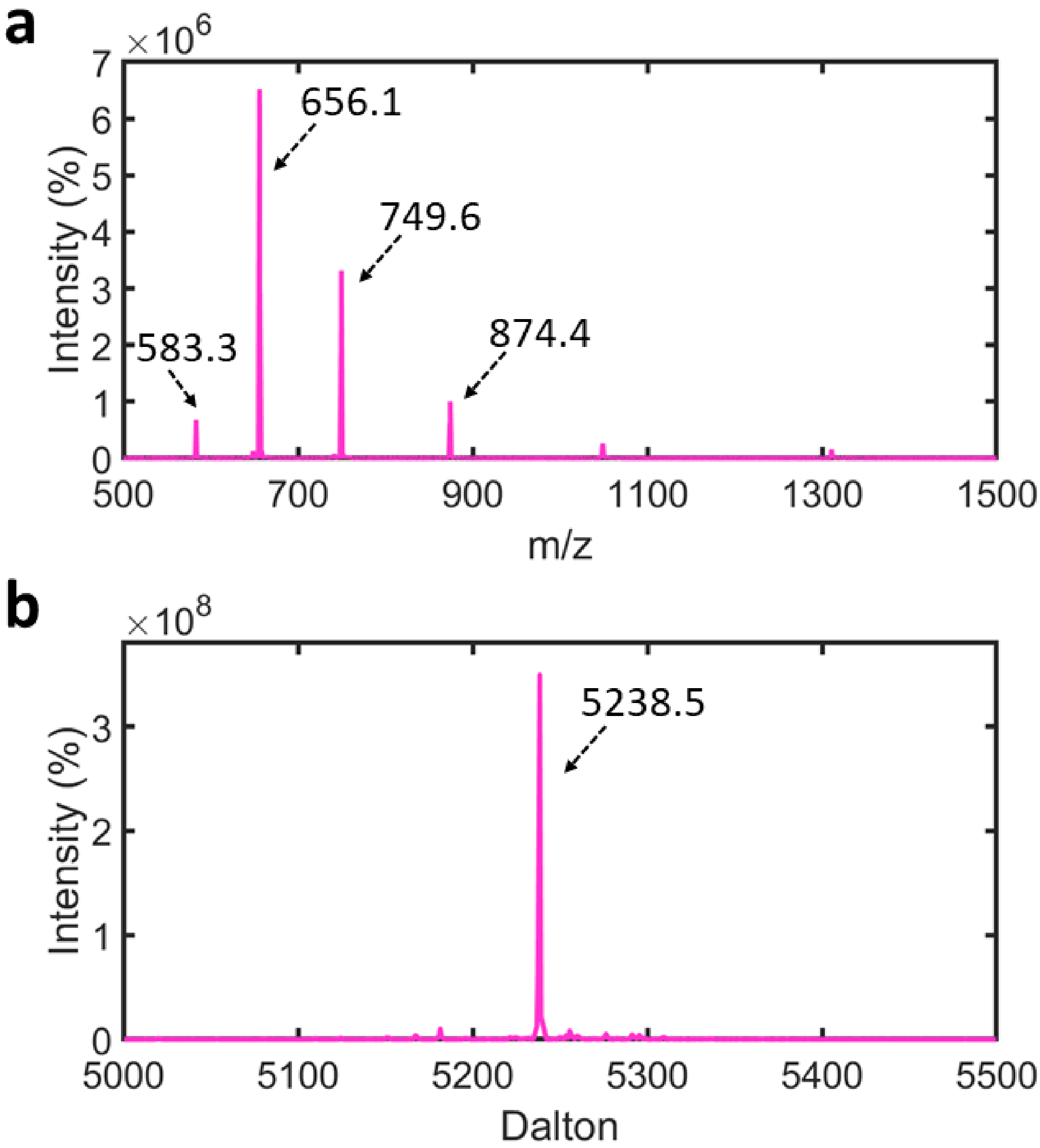


Supplementary Figure 13: **Mass analysis of TAMRA-CT45. (a)** ESI-MS spectrum TAMRA-CT45 synthesized using Fmoc-based solid-phase peptide synthesis. **(b)** The deconvoluted mass (5238.5 Da, $[M+H]^+$) corresponds with the calculated monoisotopic mass (5237.8 Da) of TAMRA-CT45.

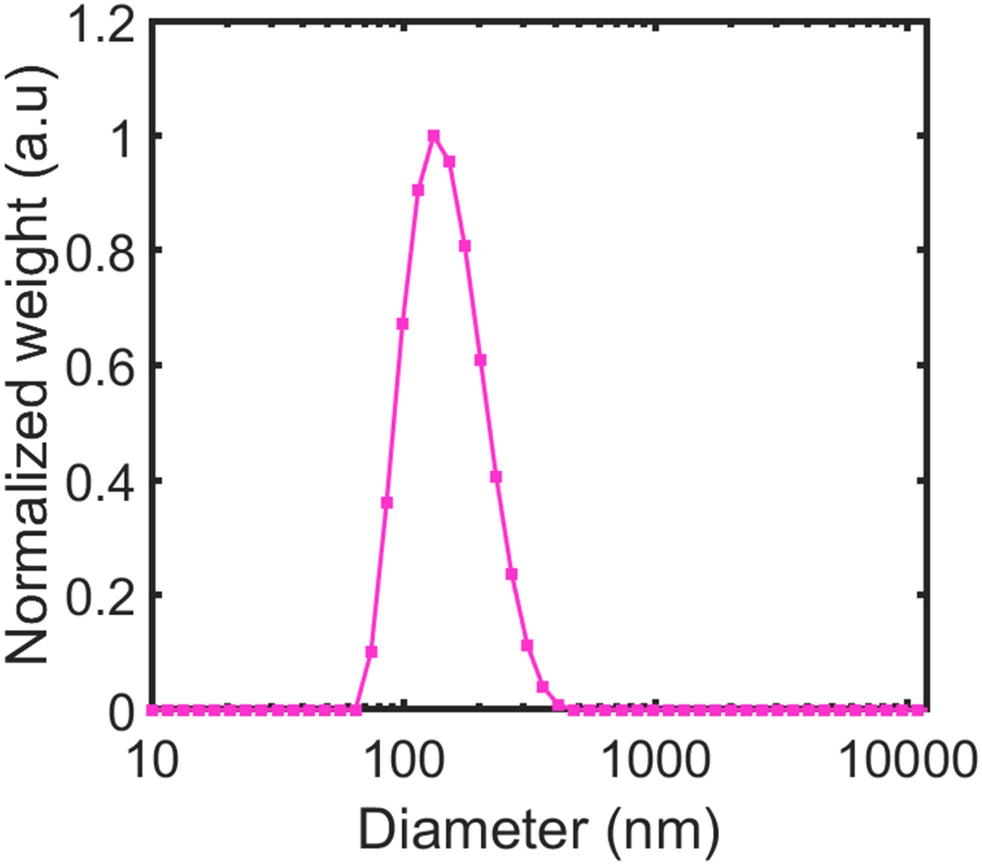


Supplementary Figure 14: **Dynamic light scattering showed the SUV diameter to be around 130 nm.** Lipid composition used was 89.9 % DOPC + 10 % DOPS + 0.1% cy5-DOPE (molar ratio) and the SUVs were produced via extrusion using a filter with a pore size of 100 nm.

#### Supplementary Video legends

##### Supporting Movie 1

**Breaking of CT shells with zebra-patterned cracks after stimulation by a hypotonic trigger.** Bright-field (left) and corresponding fluorescence (right) images showing broken CT45 shells with multiple cracks, in response to the hypotonic trigger. Some DEs burst, likely due to thinner oil shells or excessive water intake.

##### Supplementary Movie 2

**Bright-field and fluorescence $z$-stack showing the inner surface of DEs remained covered with cracks originating from broken CT45 shells.** Even 17 hours post-expansion, the cracked CT45 shell remained stable. From bottom to up, bright-field (left) images show intact DEs structure and corresponding fluorescence (right) images showing widespread CT45 shell cracks within DEs.

##### Supplementary Movie 3

**The double emulsions, along with the coated CT45 shells, shrunk upon hypertonic trigger.** Triggered by a solution consisting 4 M NaCl, bright-field (left) and corresponding fluorescence (right) images showing both DEs and CT45 shells underwent a significant shrinkage. The CT45 fluorescence intensity increased gradually, suggesting that the initial thin shells formed prior to stimulation got reinforced over time.

##### Supplementary Movie 4

**Bright-field and fluorescence $z$-stack showing reinforced CT45 shells broke at a single point after stimulation by a hypotonic trigger.** After the shrinkage-induced reinforcement by a solution consisting 1 M NaCl, CT45 shells broke with a single crack upon hypertonic trigger (salt-free OA solution). Bright-field (left) images show intact DEs and corresponding fluorescence (right) images show a single CT45 shell crack, imaged 24 hours after expansion.

**Supplementary Movie 5**

**Bright-field and fluorescence $z$-stack showing pronounced reinforced CT45 shells broken with a single crack after stimulation by a hypotonic trigger.** After the shrinkage-induced reinforcement by the solution consisting 4 M NaCl, CT45 shells broke with a single crack upon hypertonic trigger (salt-free OA solution). Bright-field (left) images show intact DEs and corresponding fluorescence (right) images show a single CT45 shell crack, imaged 24 hours after expansion.

**Supplementary Movie 6**

**CT shell formed by a rapidly evaporating acetone-water gradient was not stable.** Bright-field (left) and corresponding fluorescence (right) time-lapse images showing CT45 accumulation at the interface, shell formation, and breakage within few minutes after mixing acetone and CT45 solution on a glass slide.

**Supplementary Movie 7**

**CT45 scaffold formation and DEX partitioning within.** Bright-field (left) and CT45/DEX fluorescence (middle/right) time-lapse images showing CT45 accumulation throughout the acetone-water interface, ultimately forming a porous, skeleton-like structure. DEX fluorescence showed overlap with CT45, indicating its clear partitioning. Owing to the slow evaporation of acetone, accumulated CT45 eventually solidified and formed an interconnected network over the course of 3 hours.

#### Source legends

##### Source Data

Raw data underlying Figures 1-5 in the main article and Supplementary Figures 1c, 2, 4, 6, 12-14.
